# Structural and immunological characterization of the H3 influenza hemagglutinin during antigenic drift

**DOI:** 10.1101/2024.09.13.612776

**Authors:** Rebeca de Paiva Froes Rocha, Ilhan Tomris, Charles A. Bowman, Emma Stevens, Jason Kantorow, Weiwei Peng, Svearike Oeverdieck, James A. Ferguson, Diana D. Jung, Sander Herfst, Joost Snijder, Srirupa Chakraborty, Alba Torrents de la Peña, Zachary T. Berndsen, Robert P. de Vries, Andrew B. Ward

**Affiliations:** Department of Integrative Structural and Computational Biology, The Scripps Research Institute, La Jolla, CA 92037, USA; Brazilian Biosciences National Laboratory (LNBio), Brazilian Center for Research in Energy and Materials (CNPEM), Campinas, 13083-970, Brazil; Department of Chemical Biology and Drug Discovery, Utrecht Institute for Pharmaceutical Sciences, Utrecht University, Utrecht, The Netherlands; Department of Chemical Engineering, and Department of Chemistry and Chemical Biology, Northeastern University, Boston, MA 02115, USA; Department of Viroscience, Erasmus Medical Center, Rotterdam, The Netherlands; Biomolecular Mass Spectrometry and Proteomics, Bijvoet Center for Biomolecular Research and Utrecht Institute of Pharmaceutical Sciences, Utrecht University, Utrecht, The Netherlands; Current address: Department of Surgery, University of California San Francisco School of Medicine, San Francisco, CA 94143, USA; Current address: Department of Biochemistry, The University of Missouri, Columbia, MO, USA

**Keywords:** Influenza, antibodies, glycan evolution, hemagglutinin structure, cryoEM, vaccine design

## Abstract

The quest for a universal influenza vaccine holds great promise for mitigating the global burden of influenza-related morbidity and mortality. However, challenges persist in identifying conserved epitopes capable of inducing protection. In this study, we explore the influence of glycan evolution on H3 hemagglutinin from 1968 to present day and its impacts on antigenicity and immunogenicity. We observe that the appearance of potential N-linked glycosylation sites in Sing/16 hemagglutinin head domain reduces the binding of broadly neutralizing antibodies and shifts the polyclonal immune response upon vaccination to target the stem. Furthermore, structural characterization of HK/68 and Sing/16 by cryo-electron microscopy shows that while HK/68 is resistant to enzymatic deglycosylation, removal of glycans destabilizes the hyperglycosylated head and membrane-proximal region in Sing/16. These insights expand our understanding of glycans beyond their role in protein folding and highlight the interplay among glycan integration and immune recognition to design a universal influenza vaccine.

## Introduction

Influenza remains a significant threat to human health, with annual epidemics resulting in approximately 500,000 deaths worldwide. The hemagglutinin (HA) protein on the surface of Influenza viruses contains the major antigenic target recognized by neutralizing antibodies^1^ Genetic mutations that result in antigenic drift can render previous immunity ineffective and contribute to vaccine mismatch.^2^ Additionally, changes in the asparagine (N)-linked glycosylation pattern of HA contribute to immune evasion and have been shown to play a critical role in the emergence and spread of influenza A viruses (IAV) in humans. These include the antigenicity drop of H3N2 strain in 2003 and the emergence of the 2009 H1N1 swine flu pandemic.^3,4^

N-linked glycosylation is a critical co-translational and post-translational modification that impacts the structure and immunogenicity of viral proteins, including influenza HA.^5^ For instance, adding specific glycans during viral replication is essential for the proper folding and trafficking of HA.^6^ Glycans also shield the protein from the host immune system by preventing antibody recognition and promoting viral infectivity and replication. Thus, the accumulation of glycans in the receptor binding site (RBS) or RBS-proximal region is a strategy that the virus uses for immune evasion and to tune sialic acid receptor recognition by different glycoforms.^7^ At the same time, changes in the number and location of glycans can also expose new epitopes that can be targeted by the immune system.^8,9^ Previous studies have shown that the absence of a recently acquired glycan in a well-matched H3N2 vaccine resulted in ferrets and humans being unable to produce antibodies that efficiently neutralized circulating glycosylated viruses.^3^ Thus, the interplay between HA glycosylation and the host immune response is critical to IAV pathogenesis and strain emergence.

Influenza HA exhibits a varying number of potential N-linked glycosylation sites (PNGS) in both the variable head and the conserved stem region, dependent upon the specific virus strain and subtype. A PNGS is defined by the presence of the canonical Asn-X-Ser/Thr sequon, where X is any residue except proline. Analysis of the peptide sequence of hemagglutinins has shown that H3 circulating human strains have doubled the number of PNGS since its zoonotic emergence in 1968, with a particular concentration around the RBS.^9–12^ Importantly, the presence of a PNGS is not sufficient to guarantee glycan occupancy.^13^ While sequence analysis can be useful in predicting glycan occupancy on the HA surface, mass spectrometry-based approaches are necessary to accurately quantify and characterize the glycoforms present. Although these methods can provide valuable insights, they do not offer information on the structural changes that occur as the glycosylation on HA evolves. Furthermore, even though crystallographic studies have provided some insights into how glycans affect epitope accessibility and recognition by antibodies, these studies have not fully elucidated how glycans dynamically impact HA protein stability.^3,14^

Here, we used cryo-electron microscopy (cryo-EM), mass-spectrometry, computational modeling, and polyclonal epitope mapping to analyze how the incorporation of glycans on H3 HA from 1968 to 2022 impacts its structure, stability, antigenicity, and immunogenicity. We found that the accumulation of glycans, particularly around the highly immunogenic HA head region, shifts the overall glycan composition towards higher mannose content, directs the immune response towards the more conserved stem region, and affects the stability of the HA head domain. Hence, viral fitness and immune recognition are key determinants of shaping intrinsic HA stability and can therefore impact vaccine design and manufacturing processes.

## Results

### Emerging potential N-glycosylation sites and substitutions in H3 viruses over the years

Mutations that introduce N-linked glycosylation sites in the HA glycoprotein play a critical role in protein folding and immune evasion for human H3N2 viruses. This process may result in the addition of N-linked glycans, which can lead to substantial antigenic changes. Notably, glycan addition in H3 HAs is estimated to occur every 5 to 7 years and has been shown to impact vaccine efficacy.^3,10^ To investigate the trends in the accumulation of glycans on HA over the years, we analyzed the presence of PNGS within circulating strains spanning 1968 to 2022. We analyzed *N*-glycosidic linkages to the Asn residue of the glycosylation motif Asn-X-Ser/Thr accumulated on HA based on the analysis of ∼11,000 sequences of naturally occurring H3 strains using the code previously described by Wu et al.^7^

Figure 1A illustrates the calculated frequencies of PNGS of the total number of circulating strains each year. Raw values and percentages can be found in Supplementary Table 1. From 1968 to 2022, the number of PNGS per protomer on HA went from 7, in 1968, to 12, in 2022 (Figure 1A). PNGS such as N6 and N7, found on HA’s stem region, occur at a very low frequency, with N6 present in 1 and N7 present in 9 out of 1260 circulating strains in 2022 (Table S1). On the other hand, N8, N22, N38, N165, N285 and N483 are the most conserved and present on HA since 1968 and persist until 2022 (Figure 1A). Interestingly, these glycans all appear on esterase or stem regions of HA except for N165, which is in the globular head at the interface between protomers (Figure 1B). Other glycans that also appear on esterase or stem regions on HA such as N45, N63, N81 and N276, appear transiently (Figure 1A-B). N45 appears in less than 20% of the circulating strains and disappears between 1989 and 2010. It reappears in 30% of circulating strains in 2011, and it is found in >90% of the strains from 2014 to present (2022) (Figure 1A and Table S1). Conversely, N63, which was present in 80% of circulating strains in 1970, briefly disappears between 1971-1972, then reappears in 1973 and is still present today. N81 appears at high frequency (>50%) in 1968-1969 and 1971-1973, and completely disappears in 1993 (<10%) to present (Figure 1A and Supplementary Table 1). Similarly, N276, present in the stem region, appears in 1992 and completely disappears after 1997 (Figure 1A).

**Figure 1.**
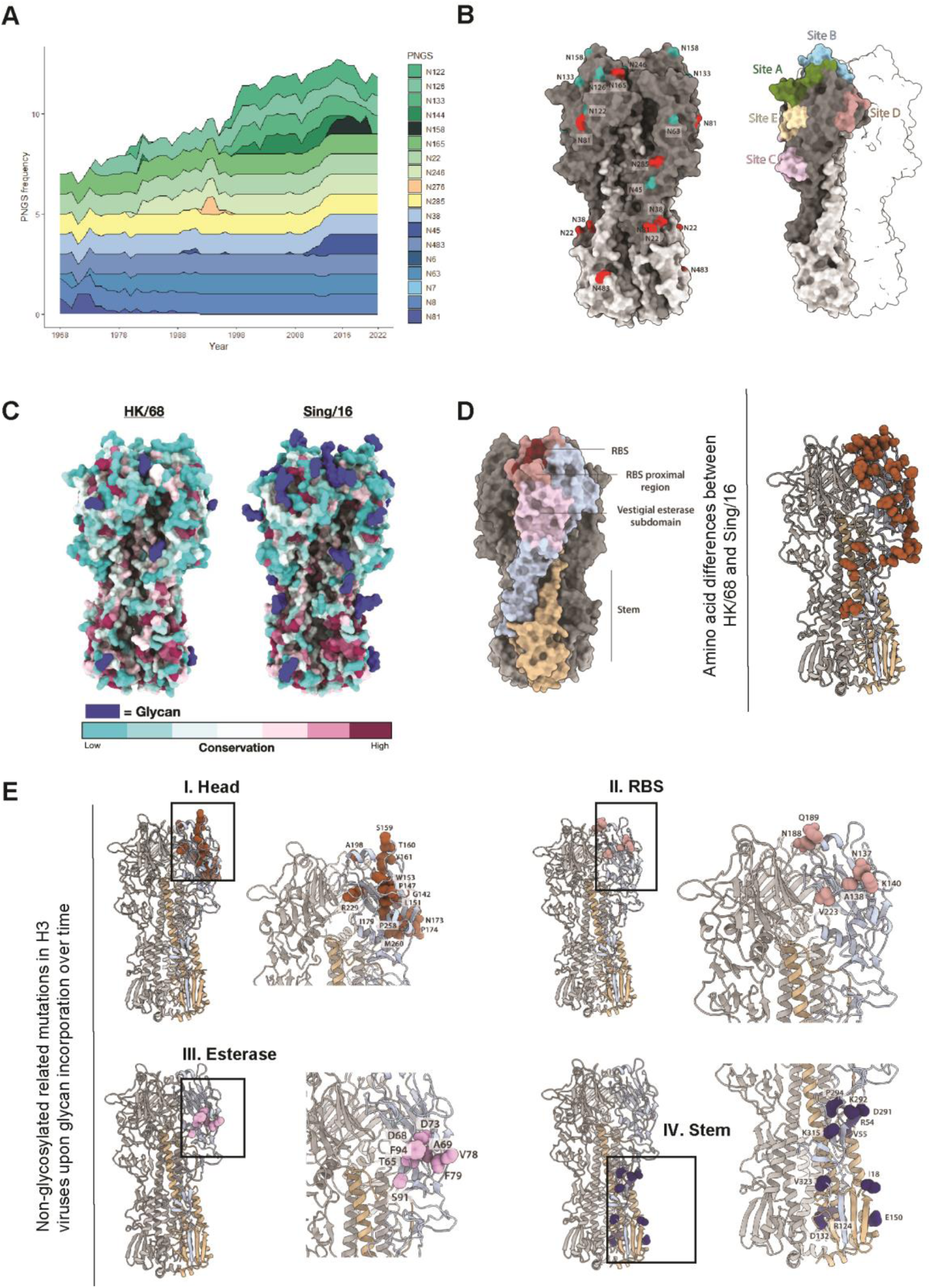
Computational analysis of N-glycosylation sites and non-glycosylation related mutations in H3 viruses over time. **A)** Stacked graph showing the frequency of each PNGS within circulating strains from 1968 to 2022. This analysis includes ∼11,000 sequences, which were downloaded from the influenza research database NCBI Virus.^22^ The Supplementary Table 1 contains the frequency of all PNGS over time. B) HA structural domains represented on HK/68 (PDB ID: 4FNK). HK/68glycans are highlighted in red and Sing/16 glycans in cyan. All residues on HA are named according to H3 consensus numbering A/Aichi/1968. C) Residue conservation of HK/68 and Sing/16 using ∼11,000 sequences from the influenza research database projected as a colormap on the HA structure. Glycans are highlighted in blue. D) HA residues with distinct amino acid identities between HK/68 and Sing/16 H3N2 strains are depicted as brown spheres on a single protomer of the HK/68 HA trimer (PDB ID: 4FNK) E) Mutations that appeared in H3 viruses at the same time as the emergence of new PNGS. The different regions of HA are highlighted in brown, salmon, pink and blue for globular head, RBS, esterase and stem regions, respectively.

By analyzing the glycans present on A/Hong Kong/1/1968 (HK/68) and comparing them to A/Singapore/INFIMH/16 (Sing/16), we observe most glycans present on HA’s globular head appear around the five canonical antigenic sites of HA: A-E (Figure 1B).^15^ Glycans such as N126, N133 and N246 appear in 1974, 1996, and 1982, respectively, and persist until 2022 (Figure 1A; Figure S1). N122, present on the RBS (Figure 1B), transiently appeared between 1980 and 1983 and then reappeared 13 years later in 1996, persisting until the present (2022) (Figure 1A). N144, present near the RBS region (Figure 1B), appeared in 1998 and completely disappeared in 2017 (Figure 1A). It then briefly reappeared in 10% of the circulating strains in 2020 only to disappear again in the following years (Figure 1A; Figure S1). N158, a key molecular determinant of antigenic distancing between influenza strains and a key residue for receptor binding^16,17^, is found in <40% of the circulating strains from 2014 to 2020 and completely disappears in the following years, like previous studies have shown.^18^ These data suggest a discernible trend regarding the emergence and accumulation of glycans on the globular head. Simultaneously, glycans appearing on the esterase and stem region tend to emerge less frequently and be more stable (Figure 1A; Figure S1). Finally, by comparing the glycans present on ∼11,000 sequences of H3, we observe that they tend to appear in non-conserved regions of Sing/16 head when compared to HK/68 (Figure 1C) and are located on the H3 head, which is consistent with studies previously performed by other groups.^10^

Next, we assessed the occurrence of non-PNGS mutations in H3 since 1968 and observed that most mutations occur within the HA head and vestigial esterase domain, while minor substitutions were observed on the esterase and stem region (Figure 1D-E). To understand where the non-PNGS mutations appeared on HA after the addition of each glycan, we examined the non-PNGS mutations in the year where the glycan was added (Figure S2). First, we observed that the globular head is the region with the highest frequency of amino acid mutations. The mutations are mostly located in loops, outside secondary structure elements (Figure 1E, panel I). Second, we detected that the amino acid mutations in the RBS are residues that are not directly involved in receptor binding (Figure 1E, panel II). This is consistent with the fact that 130-loop, 150-loop, and 190 alpha-helix, which are essential for receptor binding, are relatively conserved among HA subtypes.^7,10,19–21^

### Defining the structure and glycosylation profile of Sing/16

To better understand the complex relationship between glycan organization on the HA surface and its potential impact on emerging strains, we characterized the structures and glycosylation profile of both HK/68 and Sing/16 by cryo-EM (Figure 2 A-E and Figure S3) and mass-spectrometry (Figure 2 F-G). Notably, we obtained a structure of a fully glycosylated Sing/16 at 3.9 Å-resolution (Figure 2A and Figure S3). Both the HK/68 as the Sing/16 H3 proteins were purified from HEK293T as well as HEK293S GnTI-cells, in which N-glycans remain in a high mannose form as they lack the ability to undergo further processing into complex-type glycans. The HA plasmids contain either one or two N-terminal superfolder green fluorescent protein (sfGFP) domains, for HK/68 and Sing/16, respectively. The addition of the sfGFP increases the yield of the protein, can improve angular distribution in cryo-EM, and serve as a tool for immune tracing while not affecting antigenicity.^23^

**Figure 2.**
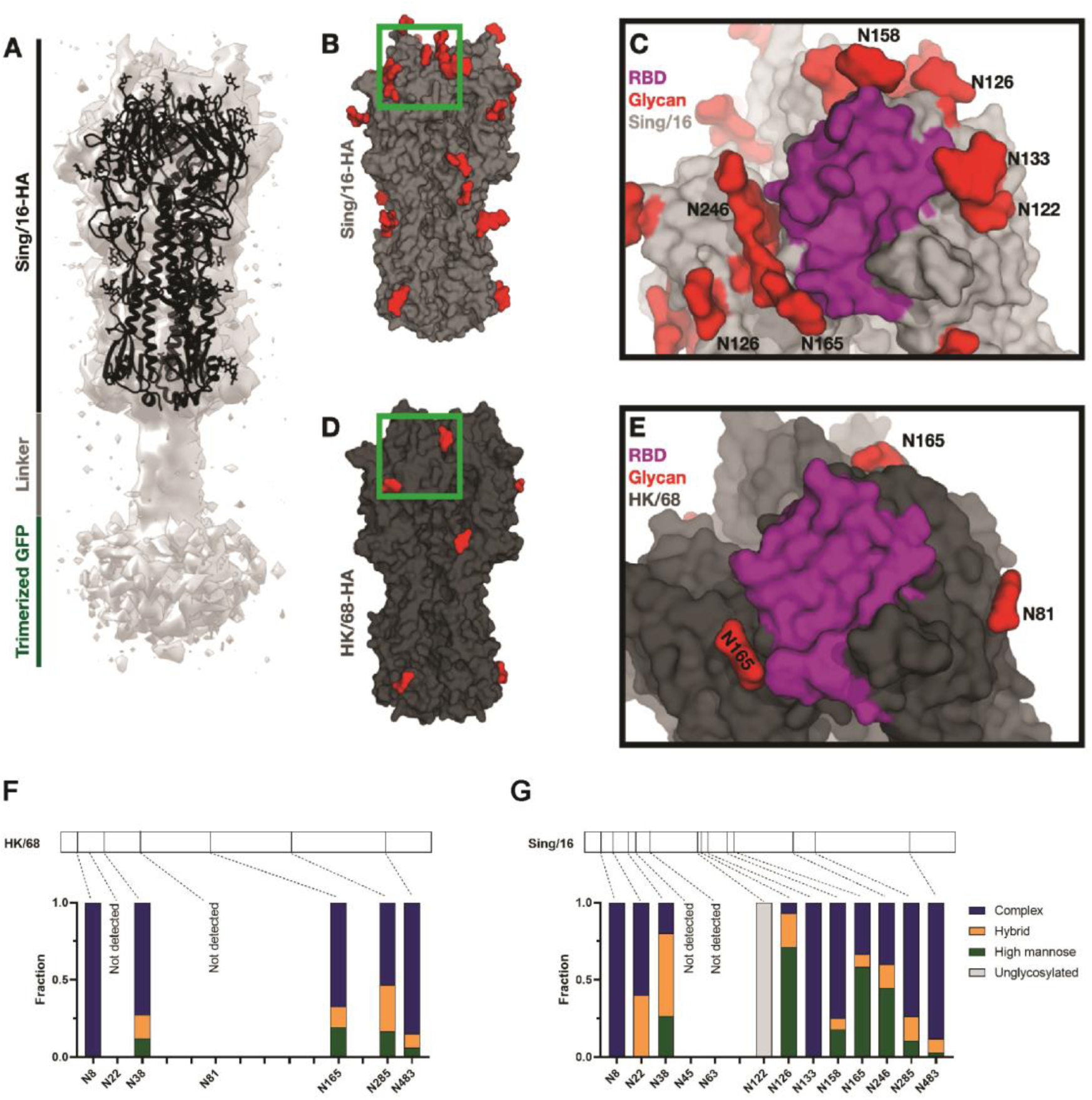
Glycan shield composition and disposition of Sing/16 and HK/86. **A)** Cryo-EM structure of trimeric A/Sing/INFIMH/16 HA. Each monomer is composed of the globular HA1 domain and the stalk-like HA2 domain. HA2 is attached to 2x sfGFP per monomer. Cryo-EM structures of Sing/16 A,D-E) and HK/68 B-C) with the glycans highlighted in red and a close-up view of the RBS region. F-G) Site-specific MS analysis of HK/68 and Sing/16 glycan shields. Bar plots show the percentage of high-mannose, complex, and unmodified peptides detected at each PNGS. At some sites, no peptides were detected. Glycan accumulation on the HA head affects epitope exposure.

The structure of Sing/16 revealed the prototypic trimeric structure of HA (Figure 2A and Figure S3). Each subunit consists of a membrane-distal polypeptide (HA1) and a membrane-proximal polypeptide (HA2). The HA1 subunit is comprised of a globular domain (residues 116 to 261) formed by an eight-stranded antiparallel β-sheet motif that includes a distal shallow pocket corresponding to the receptor-binding site, surrounded by five immunodominant epitopes (A-E; Figure 1B).^15^ The stem domain consists of a descending β-sheet motif from the HA1 subunit and is mainly formed by a helical coiled-coil structure corresponding to the HA2 subunit. We also solved the cryo-EM structure of Sing/16 produced in GnTI-cells, at 3.7Å-resolution and Sing/16 produced in HEK293T, at 3.9 Å-resolution (Figure S2). The two structures are highly similar with a Ca-root-mean-square deviation (rmsd) of 0.77Å (Figure S4), suggesting the distribution of glycan types does not have a significant impact on the folded structure. The cryo-EM structure of HK/68 was resolved to 3.4 Å-resolution, (Figure S3), and is highly similar to Sing/16 (Ca-rmsd of 0.68 Å), demonstrating that the overall structure was maintained over time despite a relatively large number of mutations and the addition of PNGS (Figure 2 B-E).

We observe density for core N-acetylglucosamine (GlcNac) moieties in our cryo-EM reconstructions (Figure 2B-E) at all PNGS on both HK/68 and Sing/16 except at N122, which is within a well-resolved region of the protein, and N8, which is near the N-terminus and is part of a poorly ordered region of the protein. Although we can observe density for individual glycans, we cannot identify the specific glycoforms and levels of occupancy from cryo-EM alone. We therefore utilized MS to determine the distribution of glycoforms at each PNGS for both strains (Figure 2F-G). While we identified glycopeptides covering most PNGS, we lack coverage at N22 and N81 on HK/68, and N45 and N63 on Sing/16, either due to low peptide detection or poor fragmentation. Of the glycans detected, however, we found that HK/68 possessed on average 76% complex, 13% hybrid, and 11% high-mannose type glycans, while Sing/16 possessed 60% complex, 17% hybrid, and 23% high-mannose. Of the PNGS that are shared between the two strains, the largest change occurred at N165, which went from 19% high-mannose on HK/68 to 58% on Sing/16, in line with previous results.^24^ N165 is located within the head domain (Figure 2C and E) where local glycan density increases, making it more difficult for processing enzymes to access.^25^ The PNGS at N122 was found to be unoccupied by any glycans on Sing/16, and while the N122 peptide was not detected from the HK/68 sample, it is also likely unoccupied given the structural conservation in this region (Figure 1C).

### Glycan accumulation on the HA head affects epitope exposure

To investigate how changes in HA glycosylation affect epitope exposure we first utilized a modified version (see Methods and Figure S7) of our previously developed high-throughput atomistic modeling pipeline to generate a large ensemble of fully glycosylated HA models of both HK/68 and Sing/16 (Figure 3A).^26,27^ From these ensembles, we calculated the “glycan encounter factor” (GEF) across the solvent-exposed surface of HA, which is a measure of the number of glycan heavy atoms encountered by an external probe approaching the protein surface (see Methods). We found that the accumulation of glycans on HA’s head domain has a dramatic effect on the epitope accessibility in the region (Figure 3B), with an average 4.7-fold increase in GEF for Sing/16 over HK/68 for head domain residues (Figure 3B and Figure S5A-B). Normalized GEF values per residue ranging from CYS52 to CYS277 were used for this calculation. In addition to the changes in shielded surface area, we also observed an increase in average inter-glycan and glycan-protein contacts of glycans on the head domain of Sing/16 compared to the same glycans present on HK/68 which is a result of increased local crowding and glycan-glycan interactions (Figure S5B). This is important because the increased local contacts indicate reduced temporal dynamics of glycans within or around antibody epitopes can have an additional impact on epitope accessibility and antibody binding beyond just steric shielding effects.^14^

**Figure 3.**
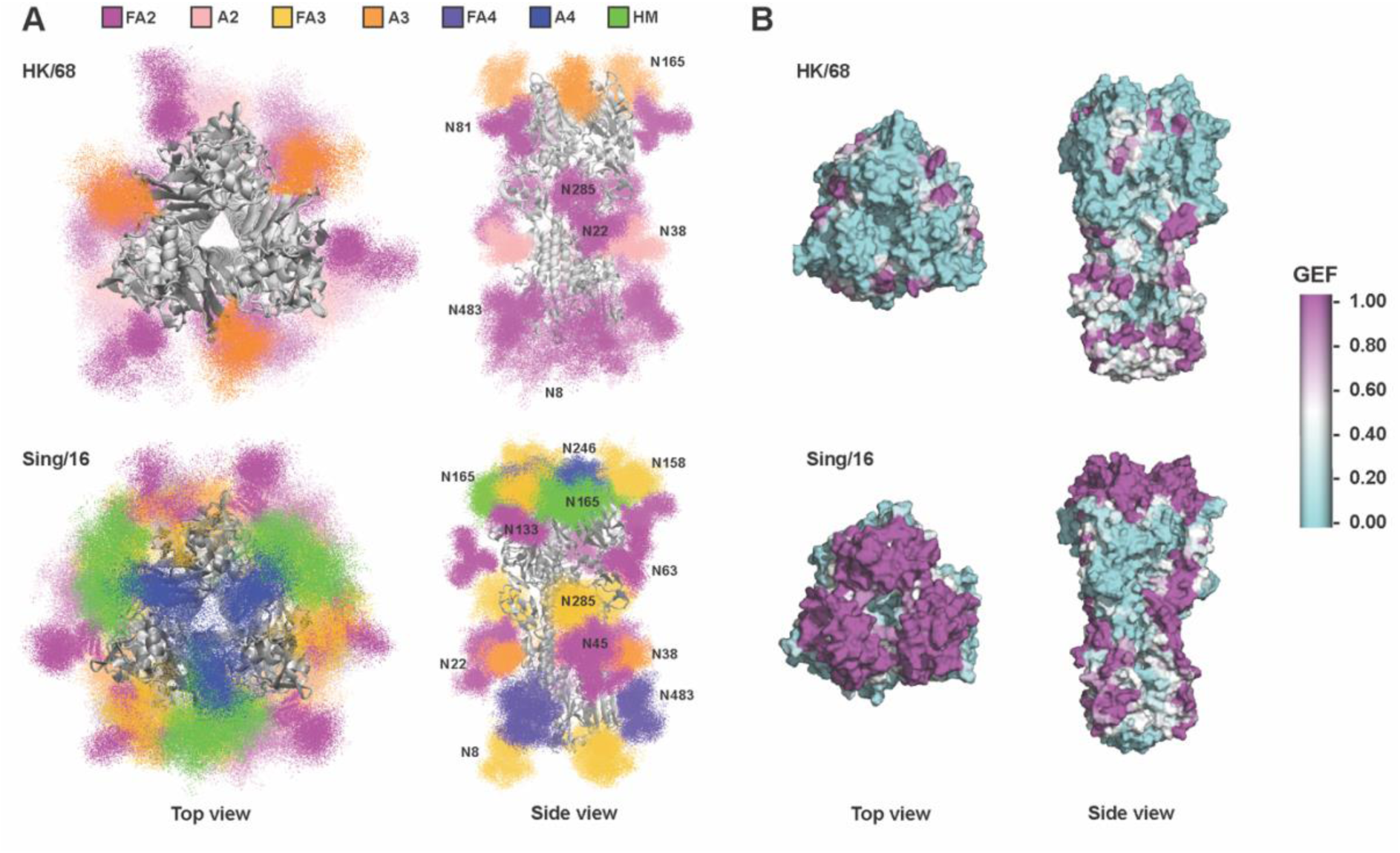
Glycan shield analysis. **A)** Models of fully glycosylated HA trimers showing 20 poses for each glycan out of the 1000 relaxed glycan conformations generated via high throughput ensemble modeling. Glycan atoms are shown as points and colored by glycan type according to the legend. Glycan types indicate whether the glycan is fucosylated with an F and the number of branching antennae. High mannose glycans are indicated by HM. One occurrence of each glycan is labeled. Images are rendered with VMD 1.9.3. **B**) Glycan encounter factor (see methods) mapped onto the surface representations of HK/68 and Sing/16. Magenta regions are considered shielded by glycans while cyan regions are accessible.

Next, we measured binding of the broadly neutralizing monoclonal antibodies (bnAbs) CR8020 and C05, which target the stem region and RBS, respectively (Figure 4) to the previously described HA variants, and found that while the glycosylation state resulting from differential expression and endo H treatment did not significantly affect the EC_50_ of the stem antibody CR8020 against HK/68 or Sing/16, it did have an impact on the binding of C05 (Figure 4A-B). Specifically, we observed overall reduced binding of C05 to Sing/16 relative to HK/68 across all conditions, suggesting the increased glycan density in the head domain partially obstructs binding. We also observed reduced binding of C05 to Sing/16 purified in HEK293S GnTI-cells relative to HEK293T, and surprisingly, even further reduction when C05 was complexed with endo H treated Sing/16, and to a lesser extent endo H treated HK/68. The C05 epitope is surrounded by numerous acquired PNGS on Sing/16 including N133, N158, N165, and N246, but it does not directly engage any of these glycans, so the reduced binding to endo H treated protein is somewhat paradoxical. However, as we will show later, de-glycosylation induced destabilization of the HA head could explain these results.

**Figure 4.**
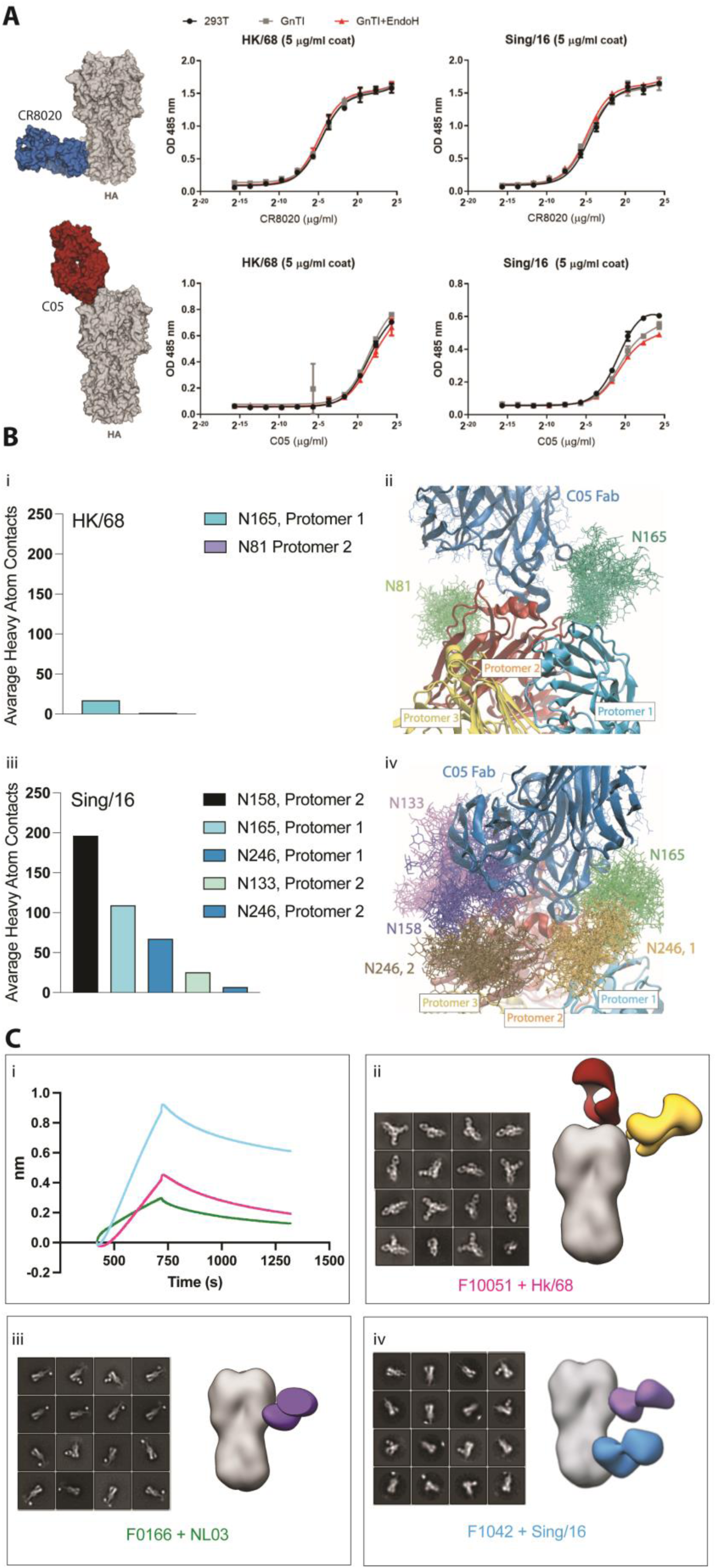
Immunogenicity analysis of HK/68 and Sing/16. **(A)** RBS-specific bnAb C05 and stem-specific bnAb CR8020 binding titers to HA measured by ELISA. HK/68 and Sing/16 HA proteins were purified from HEK 293T, HEK GnTI-cells, and HEK GnTI-cells followed by treatment with Endo H. (B) Average number of heavy atom contacts between the C05 Fab and the surrounding N-linked glycans when bound to HK/68 and Sing/16 (I, III) calculated from the computationally generated ensembles (II, IV). The threshold value used to denote a contact for both analyses was set at 4.5 Angstroms. (C) Binding of the ferret sera with the matching hemagglutinin was performed using BLI (panel i). 2D classification and side views of composite 3-D reconstructions from EMPEM analysis of polyclonal Fabs obtained from reference ferret sera (F10051, F0166 and F10421) complexed to matching protein (HK/68, NL/03 and Sing/16) purified from HEK 293T cells are shown in panels ii, iii and iv.

Finally, to evaluate how glycans impact the natural antibody response against HA, we analyzed reference sera from the Erasmus Surveillance Institute from ferrets immunized against A/Bilthoven/16190/68, A/NL/109/03 (NL/03) and Sing/16 HA by HA inhibition (HAI) assay, Biolayer Interferometry (BLI), and electron microscopy-based polyclonal epitope mapping (EMPEM).^28–30^ NL/03 was included to understand antibody responses from a time point between 1968 and 2016. For this, ferrets were inoculated with the virus intranasally, and serum was collected two weeks thereafter. HAI quantitatively determines antibody titers against HA by measuring the dilution factor of serum to inhibit the interaction between the viral HA and cell sialic acid receptor present in red blood cells through the quantification of hemagglutination.^31^ The results indicate a higher inhibition titer of the ferret antisera raised against Sing/16 (10,240) compared to HK/68 (2,560) and NL03 (160) (Table S2). Similarly, BLI analysis demonstrated a greater magnitude of binding between Sing/16 and the corresponding sera compared to HK/68 and NL/03, suggesting that ferret immunization with Sing/16 elicited a more robust immune response compared to HK/68 and NL/03 (see Figure 4C, panel I). Thus, we conducted an EMPEM assay^32^ to better understand how the polyclonal immune response elicited after immunization with the different strains differed from one another.

Our EMPEM analyses revealed that the fragment antigen binding domains (Fabs) of polyclonal antibodies complexed with matching proteins targeted the HA head in HK/68, esterase domain in NL/03, and esterase and stem domains in Sing/16 (Figure 4C). These data suggest that the increased number of glycans present on HA Sing/16 potentially favored the production of stem and esterase over head-directed antibodies against Sing/16. Further, our EMPEM data along with our BLI analysis indicate that the polyclonal stem and esterase directed antibodies against Sing/16 not only effectively neutralize the virus but also, are potentially more efficient than the head directed antibodies against HK/68.

### De-glycosylation destabilizes the HA head domain of Sing/16 but not HK/68

N-linked glycans can stabilize proteins through a variety of mechanisms,^33,34^ yet the biophysical consequences of HA glycosylation have not been thoroughly investigated. We previously showed that enzymatic deglycosylation of the Human Immunodeficiency Virus type-1 (HIV-1) Envelope glycoprotein leads to progressive destabilization and unfolding of the gp120 subunit, with the instability nucleating within the hypervariable loops 1-3 that compose the trimer apex.^26^ Considering this and the counterintuitive ELISA results showing reduced binding to the endo H-treated Sing/16, we sought to determine if HA was also being destabilized by endo H treatment. Therefore, we collected cryo-EM data of both endo H treated HK/68 and Sing/16 (Figure 5, Figure S3). Indeed, we observed significant destabilization of Sing/16 but not HK/68 following endo H treatment as seen from the cryo-EM 3-D classification results (Figure 5A). Specifically, we observed that ∼57% of the trimeric Sing/16 HA particles retained after 2-D classification exhibited some level of protein unfolding/destabilization primarily in the hyper-glycosylated head domain and to a lesser extend in the stem/membrane proximal regions (Figure 5A). By pooling the particles from the well-folded 3-D classes (100% of the HK/68 particles) we obtained 2.3 Å and 3.2 Å-resolution reconstructions of Endo H treated HK/68 and Sing/16, respectively (Figure 5B). Both structures were nearly identical to their natively glycosylated counterparts (Figure S3). Unlike Sing/16, endo H treatment of HK/68 resulted in a more uniform distribution of views resulting in a better quality and higher-resolution reconstruction (Figure S3). It is noteworthy that the resolution of 2.3 Å achieved for HK/68 represents the highest cryo-EM resolution reconstruction of HA to date, perhaps owing to the lack of glycans and isotropic tumbling in the vitreous ice on the cryo-EM grid. Difference maps calculated by subtracting the aligned Endo H treated maps from the GnT1-maps reveal the full extent of the glycosylation on both structures (Figure 5C), highlighting the nearly complete shielding of the head domain on Sing/16. Interestingly, by comparing the local map intensity around individual glycans between the GnTI- and stable endo H treated 3-D reconstructions we found that several glycans were incompletely digested by endo H (Figure 5D), most prominently the glycan at N165, which retained ∼10% and ∼50% occupancy on HK/68 and Sing/16, respectively (Figure 5E). In addition, 6 other glycans on Sing/16 also showed incomplete de-glycosylation, specifically, N8, N45, N63, N133, N246, and N285 (Figure 5E). Finally, we measured the thermal stability of both HAs before and after endo H treatment via differential scanning fluorimetry (DSF) and observed a marked reduction in melting temperature of Sing/16 relative to the fully glycosylated sample along with the appearance of a second apparent melting transition (Figure 5F).

**Figure 5.**
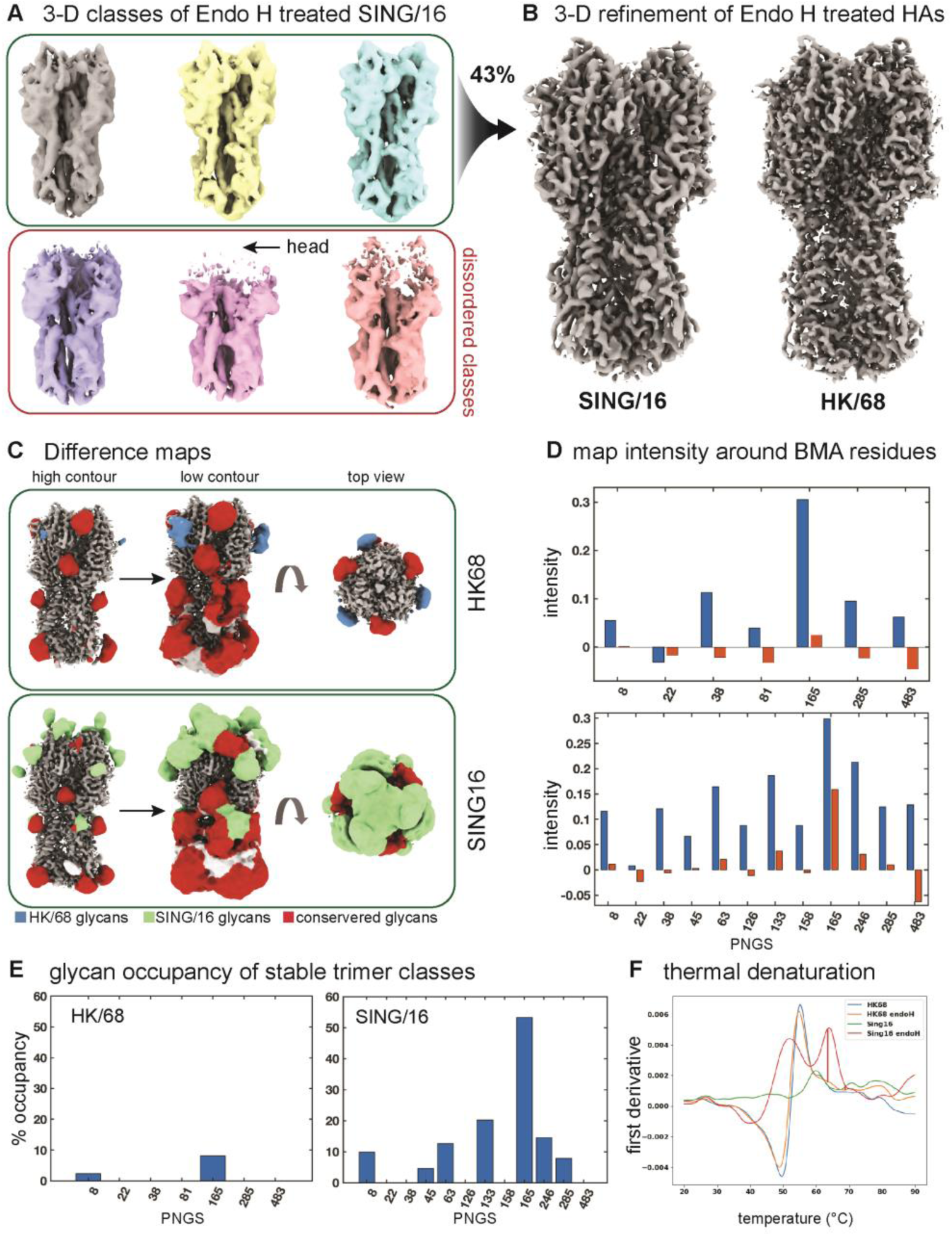
Enzymatic deglycosylation by Endo H treatment destabilizing Sing/16 HA. **A)** Cryo-EM 3-D classification results of HK/68 and Sing/16 HA after treatment with Endo H and **(B)** the resulting 3-D refinement of the selected and pooled classes. Also shown are percentages of retained and discarded particles and an overlay of 3-D reconstructions representing the two extremes of the 1^st^ principal component from 3-D variability analysis in CryoSparc. **C)** Glycan difference map of HK/68 and Sing/16 **D)** Normalized glycan mean intensity and **E)** glycan occupancy around the third glycan residue, β-mannose (BMA) between the GnTI- and stable endo H treated 3-D reconstructions of HK/68 and Sing/16 **F)** Thermostability of both HA variants obtained from nano-DSF measurements. Data was plotted using Prism 9.

## Discussion

The intricate interplay between HA glycosylation and the host immune response is a central factor influencing the pathogenesis of IAV and the emergence of viral strains. In our assessment of how the presence of glycans impacts protein heterogeneity, we observed notable changes in the dynamics of glycans within or near antibody epitopes. The increased local crowding and interactions among glycans suggest a potential mechanism influencing antibody recognition and binding (Figure 1 and 3). These changes, alongside alterations in glycan shielded surface area, can influence epitope accessibility and antibody binding, extending our understanding of glycans beyond their traditional role as steric hindrances.

Of the glycans that appeared on Sing/16 globular head, the appearance of the PNGS at N158 has been previously discussed.^12,17,35^ The complex-type fraction of the PNGS at N158 has been to be a major obstacle for α2-6 receptor binding.^16,17,36^ Interestingly, recently the NYT158-160 sequon has been replaced with NNI in currently circulating H3N2 viruses, restoring some α2-6 receptor binding. Of note, the appearance of N158 has been proposed as the most likely cause of antigenic distancing between different influenza strains circulating between 2012 and 2018.^35^

Previously published studies have proposed that the N158K substitution on influenza egg-based vaccines contributes to the loss of the glycan at N158 leading to vaccine antigenic mismatch.^18^ According to our computational analysis, PNGS at N158 appears on circulating strains between 2014 and 2020 but then disappears in 2021/2022 (Figure 1 A and Figure S1). Interestingly, the yearly CDC report highlights that, despite inadequate vaccine enrollment to generate dependable vaccine effectiveness estimates by age group or vaccine type, immunization (2021/2022) did not diminish the risk of outpatient medically attended illness associated with influenza A(H3N2) viruses.^37^ This highlights the significance of glycans when considering vaccine design and manufacturing methods.

We show that the presence of glycans on HA impacts the polyclonal antibody response in ferret specific sera as Sing/16 elicited an increased neutralizing effect when compared to HK/68. Notably, EMPEM data reveals a distinctive pattern: antibodies against HK/68 primarily targeted the head of the HA, while antibodies against Sing/16 were directed towards the esterase and stem regions (Figure 4). Interestingly, antibodies targeting the esterase domain can also inhibit hemagglutination. This is in line with our data showing that glycans did not impact the effectiveness of stem directed bnAb when comparing HK68 and Sing/16 (Figure 4). This finding aligns with the current perspective advocating for the effective use of stem-directed antibodies for effective viral neutralization. Stem-directed antibodies are increasingly recognized for their potential to provide broad-spectrum protection, exhibit reduced susceptibility to antigenic drift, and offer prospects for the development of more universal influenza vaccines.^38^

The incomplete de-glycosylation observed after endo H digestion (Figure 5D) has two implications; first, that the increased glycan density in the HA head restricts endo H access; and second, that the partially unfolded 3-D classes likely represent a subset of particles where these glycans were successfully cleaved, or conversely, that the glycans with partial occupancy in the well-folded trimer classes are particularly important for H3 stability. For instance, the glycan at N165, which shows the highest apparent occupancy in both endo H treated cryo-EM maps, is also the most ordered as measured by local map intensity (Figure 5D). It is important to note that endo H cleaves oligomannose glycans after the core GlcNac residue, thus the instability cannot be explained by the loss of local glycan-protein interactions which predominantly involve the core GlcNac.

N165 is the only conserved PNGS present on HA’s head shared between Sing/16 and HK/68. However, N165 switches from being mainly complex in HK/68 to being mainly HM in Sing/16. In fact, 3 out of 5 of the analyzed glycosides appearing on HA’s head/RBS region (N126, N165, N246) present an increased HM fraction when compared to HK/68 (Figure 2A-B). The intricate mechanisms through which clustered N-glycan sites foster high mannose species were previously discussed.^39^ Essentially, this phenomenon is believed to be driven by the presence of high density of glycans that sterically constrain the α-mannosidase-mediated glycan trimming during Golgi glycan processing. This pattern holds true across different viruses, particularly evident in HIV, where it is consistently observed across distinct clades and production systems.^40^ Finally, the differential impact of deglycosylation on Sing/16 stability relative to HK/68 may indicate that the gradual accumulation of N-linked glycans in the HA head over time have co-evolved with non-PNGS mutations that would otherwise be detrimental to HA stability and therefore viral fitness in the absence of these glycan. These deleterious mutations could be within the PNGS sequons themselves or elsewhere in the protein, such as within the more conserved domains known to be important for HA stability (Figure 1E).

In summary, our findings underscore the intricate nature of glycan-protein interactions and their significant role in modulating HA stability and immune escape both directly by blocking access to neutralizing epitopes but also allowing additional mutation of epitopes on the HA surface. Understanding the structural and biophysical interplay between HA and glycans can be used in the design of better and more stable vaccine candidates.

## Acknowledgements

We thank B. Anderson and H. L. Turner for cryo-EM data collection support and C. Bowman and J. C. Ducom for computational support. This work was supported, in part, by Bill and Melinda Gates Foundation grant INV-004923. A.T.d.l.P is a recipient of NWO Rubicon grant 45219118 and R.d.P.F.R. is supported by the Fundação de Amparo à Pesquisa do Estado de São Paulo - FAPESP Grant number: 2019/20772-4. R.P.dV is a recipient of an ERC Starting Grant from the European Commission (#802780), and a 2023 Research Grant from the Mizutani Foundation for Glycoscience. J.S and W.P. are funded by the Dutch Research Council NWO Gravitation 2013 BOO, Institute for Chemical Immunology (ICI; 024.002.009). Molecular graphics images were produced using the Chimera package from the Computer Graphics Laboratory, University of California, San Francisco (supported by NIH P41 RR-01081).

## Author contributions

R.d.P.F.R. and A.T.d.l.P. performed sequence analysis, protein purification, BLI, and NS-EM experimental procedures. Also, they did cryo-EM sample preparation, data collection and processing. Z.T.B. and S.C performed cryo-EM processing and computational analysis. J.S. performed MS experiments and analysis. S.H. provided influenza sera samples, I.T. performed ELISA, and protein expression. C.B performed computational analysis. R.P.dV provided crucial materials; R.d.P.F.R., A.T.d.l.P, S.C., Z.T.B., R.P.dV and A.B.W. wrote the paper. All authors contributed to the manuscript text by assisting in writing or providing feedback. A.T.d.l.P., R.P.dV, Z.T.B. and A.B.W. supervised the research.

## Data and materials availability

The PDB IDs and Electron Microscopy Data Bank (EMDB) IDs for the two cryo-EM structures of HK/68 (GnTI- and endo-H treated hemagglutinin) and three cryo-EM structures of Sing/16 (283F, GnTI- and endo-H treated hemagglutinin) have been deposited into the RCSB PDB (https://www.rcsb.org) under accession numbers 9CXT, 9CXU, 9D2M, 9D1U, and 9D0Y and to the EMDB database (https://www.ebi.ac.uk/emdb/) under the accession number EMD-45997, 45998, 46500, 46477, and 46466 respectively. The raw LC-MS/MS files and analyses have been deposited to the ProteomeXchange Consortium via the PRIDE partner repository with the dataset identifier PXD051492.

## Declaration of interests

The authors declare no competing interests.

## Methods

### Code glycan prevalence over time

11,187 HA amino acid sequences were pairwise aligned to reference sequence CAA24209 A/Aichi/2/1968 and analyzed using the Anchor tool package. (https://doi.org/10.1371/journal.ppat.1008753) Alignments were grouped by year, and for each year each glycan position was queried with the pattern NXT/S, representing an asparagine followed by a threonine or serine residue at position N+2. Positions 22, 23, 24, 38, 54, 61, 79, 97, 138, 142, 149, 160, 174, 181, 262, 292, 301, and 499 counted from the CAA24209 reference were queried for the year range 1968-2022 for all sequence data. Counts and totals for each matching sequence at each position were collated and analyzed for prevalence.

### Protein Expression and purification

pCD5-HA-GCN4-Fluorescent probe expression vectors were transfected into both HEK 293T and HEK293S GNTI GNT1(-) cells (which are modified HEK293S cells lacking glucosaminyl transferase I activity (ATCC CRL-3022) with polyethyleneimine I (PEI) in a 1:8 ratio (µg DNA:µg PEI) as previously described.^23^ The transfection mix was replaced after 6 hours by 293 SFM II suspension medium (Invitrogen, 11686029, supplemented with glucose 2.0 gram/L, sodium bicarbonate 3.6 gram/L, primatone 3.0 gram/L (Kerry), 1% glutaMAX (Gibco), 1.5% DMSO and 2mM valproic acid). Culture supernatants were harvested 5 days post-transfection. The HA expression was analyzed with SDS-PAGE followed by Western-blot on PVDF membrane (Biorad) using α-strep-tag mouse antibodies 1:3000 (IBA Life Sciences). Additionally, fluorescence intensities were measured using a filter based PolarStar Omega plate reader. Subsequently, HA proteins were purified with Sepharose Strep-Tactin beads (IBA Life Sciences) as previously described.

### ELISA

HK/68 and Sing/16 were HEK293T and GNT1-derived, GNT1-derived protein was further treated with Endo H (1.000 units) for three hours for the removal of high-mannose structures. Proteins were coated on maxisorp plates (Invitrogen) using at 5 µg/mL in PBS, overnight at 4 °C. Blocking was performed using 3% BSA and followed by an incubation of primary antibody CR8020 (20µg/mL, 1:4 dilution) or C05 (40µg/mL, 1:1 dilution), with an incubation of one hour at room temperature. Following the primary antibody, a secondary antibody with an HRP conjugate (1:2000) was added, with an incubation of one hour at room temperature. Hereafter the plates were developed with TMB substrate solution (34028, Thermo Scientific), the reaction was stopped after 5 minutes using 2.5 M H_2_SO_4_.

### Endoglycosidase H digestion of HK/68 and Sing/16 for cryo-EM experiments

0.5 mg of purified HK/68 or Sing/16 from HEK293S GnTI-cells was mixed with 20,000 units of Endoglycosidase H (Endo H; New England Biolabs) in non-denaturing reaction buffer to a final volume of 0.5ml and incubated at 37 °C for 5 h. To quench the reaction and purify the sample for cryoEM experiments, the HAs were run over a size exclusion column Superdex 200 increase (Cytiva life sciences) and fractions were collected and concentrated.

### Ferret samples used in the study

For the EMPEM assays used in this paper, serum samples for ferrets F10051, F0166 and F1042 obtained from Erasmus Surveillance Institute were used to test HK/68, NL/03 and Sing/16, respectively.

### BLI/OCTET

Biolayer interferometry assays were performed using the Octet Red96 instrument (Sartorius, ForteBio). IgG from serum was immobilized onto Protein A Biosensors for 120 s followed by a 60 s baseline measurement in Kinetics buffer (PBS pH 7.2 with 0.01% w/v bovine serum albumin and 0.002% v/v tween-20). The biosensors were then dipped for 300 s into wells containing HK/68, NL03 or Sing/16 diluted in kinetics buffer at a final concentration of 200nM. The sensors were dipped into wells containing kinetics buffer for 600 s to assess dissociation.

### Serum IgG isolation and Antibody digestion

IgG from ferret sera was isolated using CaptureSelect IgG-Fc Affinity Matric (Thermo Scientific) as previously described^41^. In summary, 0.5 ml of ferret sera was mixed with 0.5 ml of washed CaptureSelect resin and 4 ml of PBS. For IgG digestion, the unbound IgG was discarded, and the resin was kept. Papain was activated for 15 min at 37°C in digestion buffer (100mM Tris, 2mM EDTA, 10mM L-Cysteine, 1 mg/ml papain) and was added to the resin containing the IgG. Subsequently, digestion buffer (20 mM sodium phosphate, 10 mM EDTA, 20 mM cysteine, 0.1 mg/ml papain, pH 7.4) was added to the resin up to a total volume of 5 ml and the mixture was incubated for 4-5 h at 37°C. Iodacetamide was used to quench the reaction at a final concentration of 0.03 M and the Fab/Fc from digested IgG was purified and concentrated using the size exclusion chromatography (Superdex 200 increase column, Cytiva Life Sciences). The fractions containing purified Fabs/Fc were concentrated using 10 kDa Amicon ultrafiltration units.

### Purification of antigen-Fab complexes

Antigen-Fab complexes were generated by incubating 15 mg of antigen with 1 mg of Fab overnight at RT. The complexes were purified using a Superdex 200 increase column on Akta Pure system (GE Healthcare) running in TBS buffer. The fractions containing the complexes were concentrated using 10kDa amicon ultrafiltration units and immediately added to a negative-stain grid.

## Electron Microscopy

### Negative-stain sample preparation and imaging

Fab-antigen complexes were diluted to 20 mg/ml and applied for 10s to 400 mesh Cu grids that were carbon coated and glow discharged at 15 mA for 25 s. The Fab-antigen complex was negatively stained with 2% uranyl-formate for 50 s. Data was collected using a Tecnai Spirit electron microscope at a nominal magnification of 52,000 X with a pixel size of 2.06Å. The defocus range was set between -1.5 and -2 mm and the electron dose was 25 e^-^/Å^2^. Micrographs were recorded using a Tietz (4k) TemCam-F416 CMOS and data was acquired using the automated imaging interface from Leginon.^42^

### Negative-stain data processing

For antigen-Fab complexes, ∼150,000 particles were picked using Appion image processing package. Particles were transferred to Relion/3.0 and 2-D classification was performed.^43^ Particles that contained trimer only or trimer-Fab complexes were selected for 3-D analysis. The 3-D reference for 3-D classifications and refinements was a low-resolution model of a non-liganded HA. Initial 3-D refinement was performed using a minimum of 100,000 particles to align all the particles prior to 3-D classification. Particles were then classified into 20-40 classes and classes with similar features were combined and refined. Maps were visualized and segmented using UCSF Chimera.^44,45^

### Cryo-EM sample preparation

HK/68 and Sing/16 were diluted to a final concentration of 1 mg/ml and mixed with 0.5% Lauryl maltose neopentyl glycol (LMNG). Sing/16 was diluted to a final concentration of 7 mg/ml, 1 mg/ml and 0.2 mg/ml and mixed with 4mM CHAPSO, 0.5% LMNG and 0.3% octyl beta glucoside (OBG), when expressed in HEK293T, HEK293S GntlI and Endo H treated, respectively. UltrAufoil R 1.2/1.3 300 mesh gold grids were plasma cleaned for 20 s at 15 mA using the solarus advanced plasma cleaning system (Gatan) before loading the sample. Further, 3 μl of the sample were loaded onto the grid and plunge-frozen into nitrogen-cooled liquid ethane using the Vitrobot mark IV (Thermo Fischer Scientific). The settings for the Vitrobot were as follows: Temperature: 4°C, humidity: 100%, blotting time: 5.5 seconds, blotting force: 1 and wait time: 3.5 seconds.

### Cryo-EM data collection and image processing

Grids were loaded into a Titan Krios Cryo-Transmission Electron Microscope (FEI) operating at 300 kV and equipped with K2 Summit direct electron detector camera (Gatan). The data was collected at a total cumulative dose of 50 e^-^/Å^2^. Magnification was set at 36,000x with a resulting pixel size of 1.15 Å/pix at the specimen plane. Automated data collection was performed using Leginon software.^42^ The data collection details are presented in Table 1.

The micrograph movie frames were aligned and dose-weighted using MotionCor2.^46^ In cryoSPARC v3.1.0,^47^ patch CTF was applied and micrographs were manually curated. Further, particles were picked from micrographs using blob picker and template picker. The particles were then extracted and a few rounds of 2-D classification were performed. Particle picks were subjected to a round of 3D classification and Non-Uniform 3-D Refinement. The highest resolution classes were selected and refined in 3-D refined while imposing C3 symmetry. In Chimera, HA trimer A/Hong Kong/1/1968 (PDB entry 4FNK) crystal structure was aligned with HK/68 or Sing/16 and used as the initial model.

### Model building and refinement

The final sharpened maps were used for subsequent model building. Multiple rounds of Rosetta relax refinement^48^ and manual Coot refinement^49^ were performed. To validate the analysis, EMRinger^50^ and MolProbity^51^ were used. The final refined model was submitted to the Protein Data Bank (PDB). Structural figures were generated using UCSF Chimera X^44^ or Pymol^52^.

## Computational Modeling

### Screening and elimination of glycan burial artifacts in models

Often a molecular dynamics (MD) simulation produces a trajectory showing a glycan position that is, while thermodynamically stable, not realistically observed in experimentally derived models of the same glycoprotein.^27,53,54^ This can cause glycans to be improperly “buried” inside the protein scaffold. Since recalculating the dynamics of a glycoprotein can be time consuming, a method of quickly determining the possible burial of a glycan within the protein scaffold was needed to help discern which glycosylation site on the scaffold would, in the case of the glycan being buried, need to be structurally exposed, via homology modeling to obtain a realistic (unburied) spatial glycan position. Two separate computational methods were developed^26,27^ and tested on 3 different viral spike glycoproteins belonging to three different viruses (HIV: BG505 SOSIP SIV: gp120/gp41 complex and Human Influenza: Hemagglutinin A (HA)). Here, HIV SOSIP Envs have been used as a test system for influenza since this is an update to our previous modeling method initially designed on SOSIPs.^26,27^

Initially, it was hypothesized that measuring the Solvent Accessible Surface Area, or SASA, as a metric of the degree of glycosylation site (NGS) exposure would be a quick and computationally efficient method of predicting the possibility of glycan burial given that this site is later glycosylated without the need for the glycan to be present on the structure at the time of analysis. In theory, performing this for every NGS of each of the five viral glycoproteins would give two approximate ranges of SASA values separated by a central “cutoff” value over which an NGS’s position would or would not result in a buried glycan. A script was written in Python-3^55^ in conjunction with the Biopython^56^ module for Python-3 to parse each unglycosylated protein structure in Protein Data Bank (PDB) format. A Shrake-Rupley algorithm^57^ for calculating SASA is then performed on the structure, and the SASA value of each protein residue with respect to its adjacent residues is appended to its metadata in the structure object. Examples of the resulting SASA of each potential NGS and control non-glycosylation site asparagine residues of HA and BG505 are plotted as shown in Supplemental Figure 7 panels A and B. Although cutoff values could be approximated as the threshold between prospective buried and unburied glycans somewhat clearly in the case of HA, no correlation could be determined between SASA and glycan burial for more heavily glycosylated proteins like BG505 and gp120/gp41 as shown in Supplemental Figure 7 panel C. For this reason, a new method was developed to better classify glycan burial in a more robust, yet computationally taxing, manner.

As it was observed that most instances of glycan burial occurred at the tips of the structure and not the glycan ‘stalk’ to which the NGS is bound, a new method was developed in python to iterate over every residue in every glycan of a structure and calculate the distances from each glycan residue to the center of the protein scaffold and the carbohydrate residue’s closest surface point to the center of the protein scaffold, comparing these two values, and using this comparison to determine glycan burial; the logical details of this method are described further via the pictorial algorithmic flow chart provided in Supplemental Figure 7 panel J. The burial depth in angstroms of each sugar residue of each glycan is then plotted on a heat map for each structure to quickly determine each residue’s relative degree of burial, examples of glycan positions and their corresponding heat maps can be seen for BG505 in Supplemental Figure 7 panels D-I. Finally, an overall comparison between the entirety of the glycan shields of human influenza and HIV spike proteins can be seen in Supplemental Figure 7 panels K and L highlighting the much larger degree of glycan burial on HIV BG505. Our updated method has helped ensure that no glycans have been buried within folds of the underlying protein for the HA models used in this study.

### Modeling the Glycan Ensemble

Based on Potential N-glycosylation Site (PNGS) sequence and cryo-EM data, the glycosylation sites present on the HK/68 and SING/16 variants were determined. MS was used to select the most probable glycoform at each PNGS when possible. For glycosylation sites where the structure could not be experimentally determined, the most common glycan type for these variants was assumed (fucosylated 2-antennae, FA2). Following the methods described earlier,^26,27^ an ensemble of 3D conformations of the glycosylated hemagglutinins was generated in atomistic detail. For each hemagglutinin variant, 10 protein structures determined via cryo-EM were used as templates for glycosylation. Glycans were added to the known glycosylation sites with ideal geometries as determined by the CHARMM36 force field then randomized by 1Å across atomic coordinates. Following this randomization step, the glycan structures were relaxed via 1000 steps of conjugate gradient minimization. At this stage, the structures were equilibrated with a 500_ps molecular dynamic simulation. To ensure the conformational space was adequately sampled, five rounds of simulated annealing were performed in which the PNGS asparagine residues, loops, and glycans were unrestrained. For each of the 10 template structures, 100 glycosylated conformations were sampled. The protein backbones of these conformations were aligned to yield an ensemble of 1000 possible glycan orientations, which displays the glycan surface coverage.

### Glycan Root Mean Square Fluctuations

To demonstrate the dynamics of each glycan, the root mean square fluctuations (RMSF) at three different positions were determined. These positions were the tip (C5 of the sugar furthest from the base), the core (C5 of the β-mannose), and the base (Cα of the glycosylated asparagine). The RMSF measures the conformational fluctuations across the 1000 poses and is averaged over the three protomers present in each hemagglutinin. Calculating the RMSF of each of these positions allowed for the comparison of the fluctuations of the glycan branches and stems as well as the isolation of any backbone fluctuations

### Glycan Encounter Factor

The per-residue glycan encounter factor (GEF)^26,27^ was calculated for each variant to quantify the extent of which the glycans effectively shield the surface of the underlying protein from external probes, such as antibodies. To do so, the geometric mean of the number of heavy atoms encountered by a cylindrical probe with a 6 Å diameter approaching each residue of the proteins from the three Euclidean axes was averaged over the three protomers. A 6 Å probe was used to approximate the size of a hairpin loop – mimicking the initial encounter contact by a typical antibody. A geometric mean was performed over the x, y, z directions to ensure a residue that isaccessible from one direction, was considered unshielded. The GEF was normalized by a factor of 1.5 (approximately p = 0.05) and any values greater than one were set equal to one to indicate shielding of that residue. A GEF of zero indicates no shielding of that residue from external probes. A surface representation of the two variants was colored by the GEF to show regions of glycan coverage.

## Glycoproteomics analysis

### Glycoproteomics sample preparation

For N-linked glycan analysis, the samples were denatured at 95 °C in a final concentration of 2% sodium deoxycholate (SDC), 200 mM Tris/HCl, 10 mM tris(2-carboxyethyl)phosphine, pH 8.0 for 10 min followed with 30 min reduction at 37 °C for 30 min. Samples were next alkylated by adding 40 mM iodoacetamide and incubated in the dark at room temperature for 45 min. 3 μg sample was used for each protease digestion. Samples were split in four for parallel digestion with trypsin (Promega), chymotrypsin (Roche), alpha lytic protease (Sigma), and gluC (Sigma)-trypsin. For each protease digestion, the denatured, reduced, and alkylated samples was diluted in a total volume of 100 μL 50 mM ammonium bicarbonate, adding proteases in a 1:15 ratio (w:w) for incubation overnight at 37 °C. For the gluC-trypsin digestion, gluC was added first for two hours, followed by incubation with trypsin overnight. After overnight digestion SDC was removed through precipitation by adding 2 μL formic acid (FA) and centrifugation at 14,000g for 20 min. Following centrifugation, the supernatant containing the peptides was collected for desalting on a 30 µm Oasis HLB 96-well plate (Waters). The Oasis HLB sorbent was activated with 100% acetonitrile and subsequently equilibrated with 10% formic acid in water. Next, peptides were bound to the sorbent, washed twice with 10% formic acid in water and eluted with 100 µL of 50% acetonitrile/5% formic acid in water (v/v). The eluted peptides were vacuum-dried and resuspended in 100 µL of 2% formic acid in water.

### Glycoproteomics LC-MS/MS measurements

For each sample and protease digestion, approximately 0.15 μg of peptides were run by online reversed phase chromatography on an Agilent 1290 UHPLC coupled to a Thermo Scientific Orbitrap Fusion mass spectrometer. A Poroshell 120 EC C18 (50 cm × 75 µm, 2.7 µm, Agilent Technologies) analytical column and a ReproSil-Pur C18 (2 cm × 100 µm, 3 µm, Dr. Maisch) trap column were used for peptide separation. Samples were eluted over a 90 min gradient from 0 to 44% acetonitrile. Peptides were analyzed with a resolution setting of 60 000 in MS1. MS1 scans were obtained with an automatic gain control (AGC) target at 4e^5^, a maximum injection time of 50 ms, and a scan range of 350–2000. The precursors were selected with a 3 m/z window and fragmented by electron-transfer high-energy collision dissociation (EThcD). EThcD fragmentation was performed with calibrated charge-dependent electron-transfer dissociation (ETD) parameters and 27% NCE supplemental activation. For both fragmentation types, MS2 scans were acquired at a 30 000 resolution, a 5e^5^ AGC target, a 250 ms maximum injection time, and a scan range of 120–4000.

### Glycoproteomics data analysis

The acquired data was analyzed using Byonic (v1.4.1) against a custom database of recombinant influenza HA protein sequences and the proteases used in the experiment, searching for glycan modifications with 10/20 ppm search windows for MS1/MS2, respectively. Up to 4 missed cleavages were permitted using C-terminal cleavage at R/K for trypsin, 6 at R/K/E/D for gluC-trypsin, 6 at WFLYM for chymotrypsin or 8 at T/A/S/V for alpha lytic protease. For N-linked analysis, carbamidomethylation of cysteine was set as fixed modification, oxidation of methionine/tryptophan as variable rare 2. N-glycan modifications were set as variable common 2, allowing up to max. 2 variable common and 2 rare modification per peptide. All N-linked glycan databases from Byonic were merged into a single non-redundant list to be included in the database search. All reported glycopeptides in the Byonic result files were manually inspected for quality of fragment assignments (with scores ≥ 200). Glycans were classified based on HexNAc content as truncated (≤ 2 HexNAc; < 3 Hex), paucimannose (2 HexNAc, 3 Hex), high mannose (2 HexNAc; > 3 Hex), hybrid (3 HexNAc) or complex (> 3 HexNAc). Reported peptide spectral matches (PSMs) were pooled based on the number of HexNAc, Fuc or NeuAc residues to distinguish truncated, paucimannose, high mannose, hybrid, and complex glycosylation, or the degree of fucosylation and sialylation, respectively.

## Notes

### Competing Interest Statement

The authors have declared no competing interest.

## References

1. About Estimated Flu Burden | Flu Burden | CDC https://www.cdc.gov/flu-burden/php/about/index.html?CDC_AAref_Val=https://www.cdc.gov/flu/about/burden/index.html.

2. Kim, H., Webster, R.G., and Webby, R.J. (2018). Influenza Virus: Dealing with a Drifting and Shifting Pathogen. Viral Immunol. 31, 174–183. 10.1089/vim.2017.0141.

3. Wu, N.C., Zost, S.J., Thompson, A.J., Oyen, D., Nycholat, C.M., McBride, R., Paulson, J.C., Hensley, S.E., and Wilson, I.A. (2017). A structural explanation for the low effectiveness of the seasonal influenza H3N2 vaccine. PLoS Pathog. 13, e1006682. 10.1371/journal.ppat.1006682.

4. Maines, T.R., Jayaraman, A., Belser, J.A., Wadford, D.A., Pappas, C., Zeng, H., Gustin, K.M., Pearce, M.B., Viswanathan, K., Shriver, Z.H., et al. (2009). Transmission and Pathogenesis of Swine-Origin 2009 A(H1N1) Influenza Viruses in Ferrets and Mice. Science 325, 484–487. 10.1126/science.1177238.

5. Shrimal, S., Cherepanova, N.A., and Gilmore, R. (2015). Cotranslational and posttranslocational N-glycosylation of proteins in the endoplasmic reticulum. Semin. Cell Dev. Biol. 41, 71–78. 10.1016/j.semcdb.2014.11.005.

6. Daniels, R., Kurowski, B., Johnson, A.E., and Hebert, D.N. (2003). N-Linked Glycans Direct the Cotranslational Folding Pathway of Influenza Hemagglutinin. Mol. Cell 11, 79–90. 10.1016/s1097-2765(02)00821-3.

7. Wu, N.C., and Wilson, I.A. (2017). A Perspective on the Structural and Functional Constraints for Immune Evasion: Insights from Influenza Virus. J Mol Biol 429, 2694–2709. 10.1016/j.jmb.2017.06.015.

8. Boyoglu-Barnum, S., Hutchinson, G.B., Boyington, J.C., Moin, S.M., Gillespie, R.A., Tsybovsky, Y., Stephens, T., Vaile, J.R., Lederhofer, J., Corbett, K.S., et al. (2020). Glycan repositioning of influenza hemagglutinin stem facilitates the elicitation of protective cross-group antibody responses. Nat. Commun. 11, 791. 10.1038/s41467-020-14579-4.

9. Bajic, G., Maron, M.J., Adachi, Y., Onodera, T., McCarthy, K.R., McGee, C.E., Sempowski, G.D., Takahashi, Y., Kelsoe, G., Kuraoka, M., et al. (2019). Influenza Antigen Engineering Focuses Immune Responses to a Subdominant but Broadly Protective Viral Epitope. Cell Host Microbe 25, 827–835.e6. 10.1016/j.chom.2019.04.003.

10. Altman, M.O., Angel, M., Košík, I., Trovão, N.S., Zost, S.J., Gibbs, J.S., Casalino, L., Amaro, R.E., Hensley, S.E., Nelson, M.I., et al. (2019). Human Influenza A Virus Hemagglutinin Glycan Evolution Follows a Temporal Pattern to a Glycan Limit. mBio 10, 10.1128/mbio.00204-19. https://doi.org/10.1128/mbio.00204-19.

11. Alymova, I.V., York, I.A., Air, G.M., Cipollo, J.F., Gulati, S., Baranovich, T., Kumar, A., Zeng, H., Gansebom, S., and McCullers, J.A. (2016). Glycosylation changes in the globular head of H3N2 influenza hemagglutinin modulate receptor binding without affecting virus virulence. Sci. Rep. 6, 36216. 10.1038/srep36216.

12. Thompson, A.J., Cao, L., Ma, Y., Wang, X., Diedrich, J.K., Kikuchi, C., Willis, S., Worth, C., McBride, R., Yates, J.R., et al. (2020). Human Influenza Virus Hemagglutinins Contain Conserved Oligomannose N-Linked Glycans Allowing Potent Neutralization by Lectins. Cell Host Microbe 27, 725–735.e5. 10.1016/j.chom.2020.03.009.

13. Chuang, G.-Y., Boyington, J.C., Joyce, M.G., Zhu, J., Nabel, G.J., Kwong, P.D., and Georgiev, I. (2012). Computational prediction of N-linked glycosylation incorporating structural properties and patterns. Bioinformatics 28, 2249–2255. 10.1093/bioinformatics/bts426.

14. Wang, C.-C., Chen, J.-R., Tseng, Y.-C., Hsu, C.-H., Hung, Y.-F., Chen, S.-W., Chen, C.-M., Khoo, K.-H., Cheng, T.-J., Cheng, Y.-S.E., et al. (2009). Glycans on influenza hemagglutinin affect receptor binding and immune response. Proc. Natl. Acad. Sci. 106, 18137–18142. 10.1073/pnas.0909696106.

15. Wu, N.C., and Wilson, I.A. (2020). Structural Biology of Influenza Hemagglutinin: An Amaranthine Adventure. Viruses 12, 1053. 10.3390/v12091053.

16. Thompson, A.J., Wu, N.C., Canales, A., Kikuchi, C., Zhu, X., Toro, B.F. de, Cañada, F.J., Worth, C., Wang, S., McBride, R., et al. (2024). Evolution of human H3N2 influenza virus receptor specificity has substantially expanded the receptor-binding domain site. Cell Host Microbe 32, 261–275.e4. 10.1016/j.chom.2024.01.003.

17. Unione, L., Ammerlaan, A.N.A., Bosman, G.P., Uslu, E., Liang, R., Broszeit, F., Woude, R. van der Liu, Y., Ma, S., Liu, L., et al. (2024). Probing altered receptor specificities of antigenically drifting human H3N2 viruses by chemoenzymatic synthesis, NMR, and modeling. Nat. Commun. 15, 2979. 10.1038/s41467-024-47344-y.

18. Bolton, M.J., Ort, J.T., McBride, R., Swanson, N.J., Wilson, J., Awofolaju, M., Furey, C., Greenplate, A.R., Drapeau, E.M., Pekosz, A., et al. (2022). Antigenic and virological properties of an H3N2 variant that continues to dominate the 2021–22 Northern Hemisphere influenza season. Cell Rep. 39, 110897. 10.1016/j.celrep.2022.110897.

19. Martin, D.J., and Schoub, B.D. (1997). Influenza--the forgotten vaccination. S. Afr. Méd. J. Suid-Afr. Tydskr. vir Geneeskd. 87, 869–871.

20. Yang, D., Liu, J., Ju, H., Ge, F., Wang, J., Li, X., Zhou, J., and Liu, P. (2015). Genetic analysis of H3N2 avian influenza viruses isolated from live poultry markets and poultry slaughterhouses in Shanghai, China in 2013. Virus Genes 51, 25–32. 10.1007/s11262-015-1198-5.

21. Wu, N.C., Xie, J., Zheng, T., Nycholat, C.M., Grande, G., Paulson, J.C., Lerner, R.A., and Wilson, I.A. (2017). Diversity of Functionally Permissive Sequences in the Receptor-Binding Site of Influenza Hemagglutinin. Cell Host Microbe 21, 742–753.e8. 10.1016/j.chom.2017.05.011.

22. NCBI Virus https://www.ncbi.nlm.nih.gov/labs/virus/vssi/#/.

23. Nemanichvili, N., Tomris, I., Turner, H.L., McBride, R., Grant, O.C., Woude, R. van der Aldosari, M.H., Pieters, R.J., Woods, R.J., Paulson, J.C., et al. (2019). Fluorescent Trimeric Hemagglutinins Reveal Multivalent Receptor Binding Properties. J Mol Biol 431, 842–856. 10.1016/j.jmb.2018.12.014.

24. Lee, C.-C.D., Watanabe, Y., Wu, N.C., Han, J., Kumar, S., Pholcharee, T., Seabright, G.E., Allen, J.D., Lin, C.-W., Yang, J.-R., et al. (2021). A cross-neutralizing antibody between HIV-1 and influenza virus. Plos Pathog 17, e1009407. 10.1371/journal.ppat.1009407.

25. Pritchard, L.K., Vasiljevic, S., Ozorowski, G., Seabright, G.E., Cupo, A., Ringe, R., Kim, H.J., Sanders, R.W., Doores, K.J., Burton, D.R., et al. (2015). Structural Constraints Determine the Glycosylation of HIV-1 Envelope Trimers. Cell Reports 11, 1604–1613. 10.1016/j.celrep.2015.05.017.

26. Berndsen, Z.T., Chakraborty, S., Wang, X., Cottrell, C.A., Torres, J.L., Diedrich, J.K., López, C.A., Yates, J.R., Gils, M.J. van, Paulson, J.C., et al. (2020). Visualization of the HIV-1 Env glycan shield across scales. P Natl Acad Sci Usa 117, 28014–28025. 10.1073/pnas.2000260117.

27. Chakraborty, S., Berndsen, Z.T., Hengartner, N.W., Korber, B.T., Ward, A.B., and Gnanakaran, S. (2020). Quantification of the Resilience and Vulnerability of HIV-1 Native Glycan Shield at Atomistic Detail. Iscience 23, 101836. 10.1016/j.isci.2020.101836.

28. Bianchi, M., Turner, H.L., Nogal, B., Cottrell, C.A., Oyen, D., Pauthner, M., Bastidas, R., Nedellec, R., McCoy, L.E., Wilson, I.A., et al. (2018). Electron-Microscopy-Based Epitope Mapping Defines Specificities of Polyclonal Antibodies Elicited during HIV-1 BG505 Envelope Trimer Immunization. Immunity 49, 288–300.e8. 10.1016/j.immuni.2018.07.009.

29. Han, J., Schmitz, A.J., Richey, S.T., Dai, Y.-N., Turner, H.L., Mohammed, B.M., Fremont, D.H., Ellebedy, A.H., and Ward, A.B. (2021). Polyclonal epitope mapping reveals temporal dynamics and diversity of human antibody responses to H5N1 vaccination. Cell Rep. 34, 108682. 10.1016/j.celrep.2020.108682.

30. Turner, H.L., Andrabi, R., Cottrell, C.A., Richey, S.T., Song, G., Callaghan, S., Anzanello, F., Moyer, T.J., Abraham, W., Melo, M., et al. (2021). Disassembly of HIV envelope glycoprotein trimer immunogens is driven by antibodies elicited via immunization. Sci Adv 7, eabh2791. 10.1126/sciadv.abh2791.

31. Medeiros, R., Escriou, N., Naffakh, N., Manuguerra, J.-C., and Werf, S. van der (2001). Hemagglutinin Residues of Recent Human A(H3N2) Influenza Viruses That Contribute to the Inability to Agglutinate Chicken Erythrocytes. Virology 289, 74–85. 10.1006/viro.2001.1121.

32. Turner, H.L., Jackson, A.M., Richey, S.T., Sewall, L.M., Antanasijevic, A., Hangartner, L., and Ward, A.B. (2023). Protocol for analyzing antibody responses to glycoprotein antigens using electron-microscopy-based polyclonal epitope mapping. STAR Protoc. 4, 102476. 10.1016/j.xpro.2023.102476.

33. Zhou, Q., and Qiu, H. (2019). The Mechanistic Impact of N-Glycosylation on Stability, Pharmacokinetics, and Immunogenicity of Therapeutic Proteins. J. Pharm. Sci. 108, 1366– 1377. 10.1016/j.xphs.2018.11.029.

34. Tan, N.Y., Bailey, U.-M., Jamaluddin, M.F., Mahmud, S.H.B., Raman, S.C., and Schulz, B.L. (2014). Sequence-based protein stabilization in the absence of glycosylation. Nat. Commun. 5, 3099. 10.1038/ncomms4099.

35. Jorquera, P.A., Mishin, V.P., Chesnokov, A., Nguyen, H.T., Mann, B., Garten, R., Barnes, J., Hodges, E., Cruz, J.D.L., Xu, X., et al. (2019). Insights into the antigenic advancement of influenza A(H3N2) viruses, 2011–2018. Sci. Rep. 9, 2676. 10.1038/s41598-019-39276-1.

36. Ji, Y., White, Y.J., Hadden, J.A., Grant, O.C., and Woods, R.J. (2017). New insights into influenza A specificity: an evolution of paradigms. Curr. Opin. Struct. Biol. 44, 219–231. 10.1016/j.sbi.2017.06.001.

37. Chung, J.R., Kim, S.S., Kondor, R.J., Smith, C., Budd, A.P., Tartof, S.Y., Florea, A., Talbot, H.K., Grijalva, C.G., Wernli, K.J., et al. (2022). Interim Estimates of 2021–22 Seasonal Influenza Vaccine Effectiveness — United States, February 2022. Morb. Mortal. Wkly. Rep. 71, 365–370. 10.15585/mmwr.mm7110a1.

38. Neerukonda, S.N., Vassell, R., and Weiss, C.D. (2020). Neutralizing Antibodies Targeting the Conserved Stem Region of Influenza Hemagglutinin. Vaccines 8, 382. 10.3390/vaccines8030382.

39. Behrens, A.-J., and Crispin, M. (2017). Structural principles controlling HIV envelope glycosylation. Curr. Opin. Struct. Biol. 44, 125–133. 10.1016/j.sbi.2017.03.008.

40. Bonomelli, C., Doores, K.J., Dunlop, D.C., Thaney, V., Dwek, R.A., Burton, D.R., Crispin, M., and Scanlan, C.N. (2011). The Glycan Shield of HIV Is Predominantly Oligomannose Independently of Production System or Viral Clade. PLoS ONE 6, e23521. 10.1371/journal.pone.0023521.

41. Peña, A.T. de la, Sewall, L.M., Rocha, R. de P.F., Jackson, A.M., Pratap, P.P., Bangaru, S., Cottrell, C.A., Mohanty, S., Shaw, A.C., and Ward, A.B. (2023). Increasing sensitivity of antibody-antigen interactions using photo-cross-linking. Cell Rep. Methods 3, 100509. 10.1016/j.crmeth.2023.100509.

42. Suloway, C., Pulokas, J., Fellmann, D., Cheng, A., Guerra, F., Quispe, J., Stagg, S., Potter, C.S., and Carragher, B. (2005). Automated molecular microscopy: The new Leginon system. J Struct Biol 151, 41–60. 10.1016/j.jsb.2005.03.010.

43. Scheres, S.H.W. (2012). RELION: Implementation of a Bayesian approach to cryo-EM structure determination. J. Struct. Biol. 180, 519–530. 10.1016/j.jsb.2012.09.006.

44. Pettersen, E.F., Goddard, T.D., Huang, C.C., Meng, E.C., Couch, G.S., Croll, T.I., Morris, J.H., and Ferrin, T.E. (2021). UCSF ChimeraX: Structure visualization for researchers, educators, and developers. Protein Sci 30, 70–82. 10.1002/pro.3943.

45. Pettersen, E.F., Goddard, T.D., Huang, C.C., Couch, G.S., Greenblatt, D.M., Meng, E.C., and Ferrin, T.E. (2004). UCSF Chimera—A visualization system for exploratory research and analysis. J Comput Chem 25, 1605–1612. 10.1002/jcc.20084.

46. Zheng, S.Q., Palovcak, E., Armache, J.-P., Verba, K.A., Cheng, Y., and Agard, D.A. (2017). MotionCor2: anisotropic correction of beam-induced motion for improved cryo-electron microscopy. Nat Methods 14, 331–332. 10.1038/nmeth.4193.

47. Punjani, A., Rubinstein, J.L., Fleet, D.J., and Brubaker, M.A. (2017). cryoSPARC: algorithms for rapid unsupervised cryo-EM structure determination. Nat. Methods 14, 290–296. 10.1038/nmeth.4169.

48. Wang, R.Y.-R., Song, Y., Barad, B.A., Cheng, Y., Fraser, J.S., and DiMaio, F. (2016). Automated structure refinement of macromolecular assemblies from cryo-EM maps using Rosetta. Elife 5, e17219. 10.7554/elife.17219.

49. Emsley, P., and Crispin, M. (2018). Structural analysis of glycoproteins: building N-linked glycans with Coot. Acta Crystallogr Sect D Struct Biology 74, 256–263. 10.1107/s2059798318005119.

50. Barad, B.A., Echols, N., Wang, R.Y.-R., Cheng, Y., DiMaio, F., Adams, P.D., and Fraser, J.S. (2015). EMRinger: Side-chain-directed model and map validation for 3D Electron Cryomicroscopy. Nat Methods 12, 943–946. 10.1038/nmeth.3541.

51. Williams, C.J., Headd, J.J., Moriarty, N.W., Prisant, M.G., Videau, L.L., Deis, L.N., Verma, V., Keedy, D.A., Hintze, B.J., Chen, V.B., et al. (2018). MolProbity: More and better reference data for improved all-atom structure validation. Protein Sci 27, 293–315. 10.1002/pro.3330.

52. Schrodinger, and LLC (2015). The PyMOL Molecular Graphics System, Version 3.0.

53. Chakraborty, S., Wagh, K., Gnanakaran, S., and López, C.A. (2021). Development of Martini 2.2 parameters for N-glycans: a case study of the HIV-1 Env glycoprotein dynamics. Glycobiology 31, 787–799. 10.1093/glycob/cwab017.

54. Watanabe, Y., Berndsen, Z.T., Raghwani, J., Seabright, G.E., Allen, J.D., Pybus, O.G., McLellan, J.S., Wilson, I.A., Bowden, T.A., Ward, A.B., et al. (2020). Vulnerabilities in coronavirus glycan shields despite extensive glycosylation. Nat Commun 11, 2688. 10.1038/s41467-020-16567-0.

55. The Python Language Reference — Python 3.12.5 documentation https://docs.python.org/3/reference/index.html.

56. Cock, P.J.A., Antao, T., Chang, J.T., Chapman, B.A., Cox, C.J., Dalke, A., Friedberg, I., Hamelryck, T., Kauff, F., Wilczynski, B., et al. (2009). Biopython: freely available Python tools for computational molecular biology and bioinformatics. Bioinformatics 25, 1422– 1423. 10.1093/bioinformatics/btp163.

57. Shrake, A., and Rupley, J.A. (1973). Environment and exposure to solvent of protein atoms. Lysozyme and insulin. J. Mol. Biol. 79, 351–371. 10.1016/0022-2836(73)90011-9.

