## Supplemental Material for "Structural and immunological characterization of the H3 influenza hemagglutinin during antigenic drift"

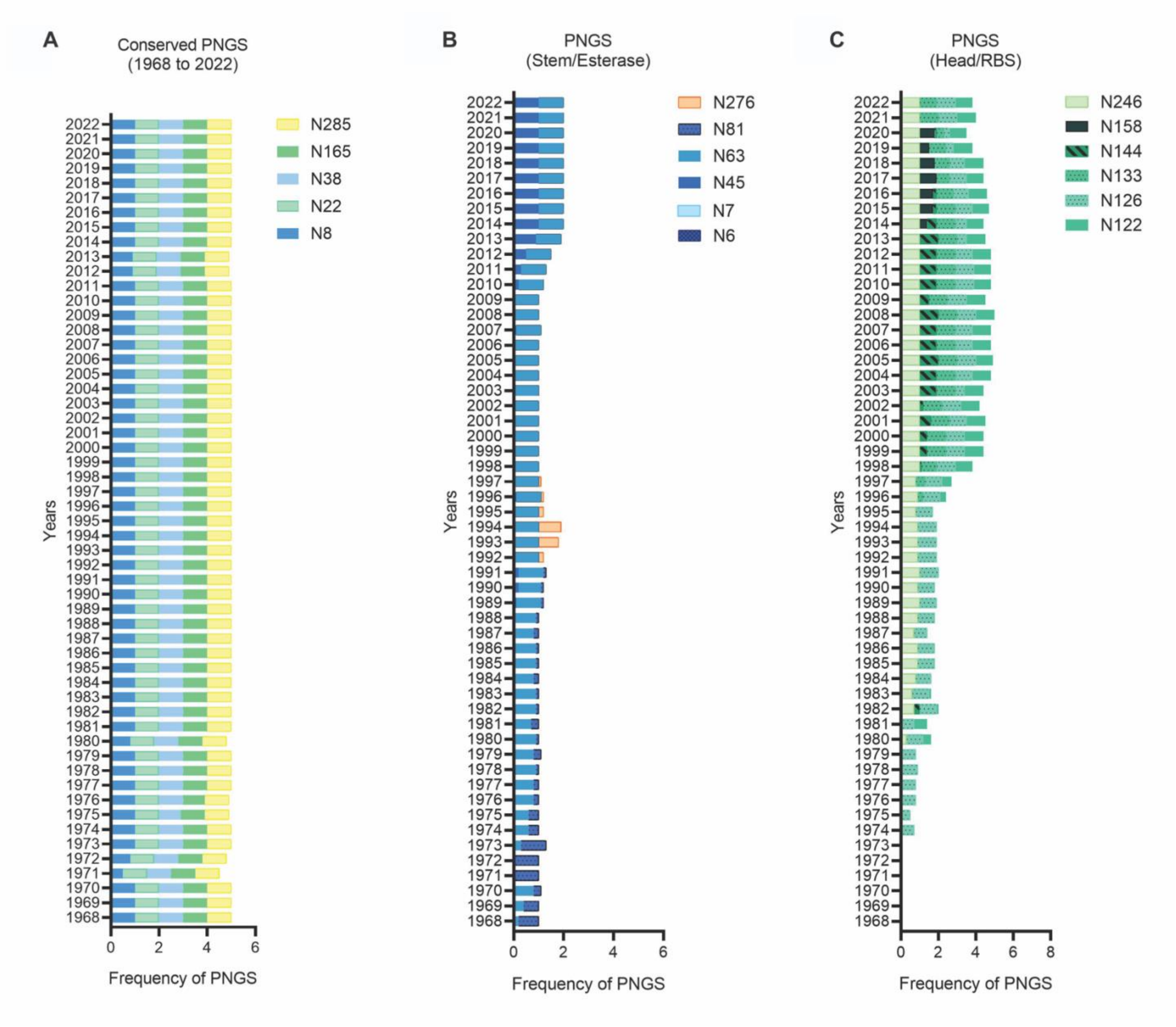

**Supplementary Figure 1. Evolution of PNGS distribution in different regions of HA over time.** (**A**) Glycans that are consistently present on H3 circulating strains from 1968 to 2022. (**B**) Glycans present on stem and esterase regions and (**C**) glycans present on head and RBS regions of H3 circulating strains from 1968 to 2022. This analysis includes ~11000 sequences, which were downloaded from the influenza research database.[22]

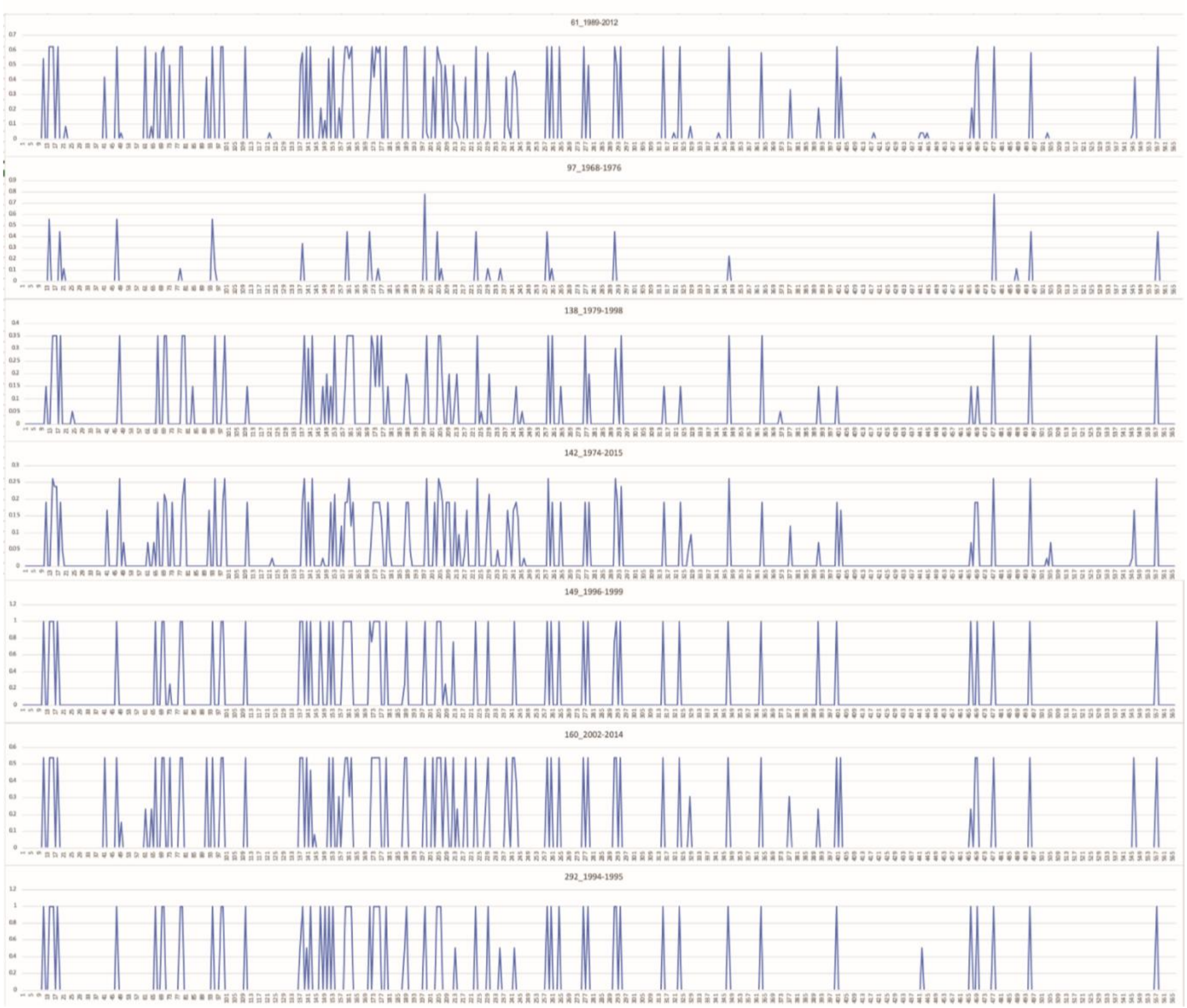

**Supplementary Figure 2. Non-PNGS mutations on H3 HA circulating strains over time.** Analysis of the non-glycosylation related mutations on H3 HA circulating strains on the year interval when the glycan was added. The non-glycosylation related mutations on HA H3 circulating strains were determined after the analysis of ~11,000 sequences, which were downloaded from the influenza research database.[22] The numbering corresponds to HK/68 reference sequence.

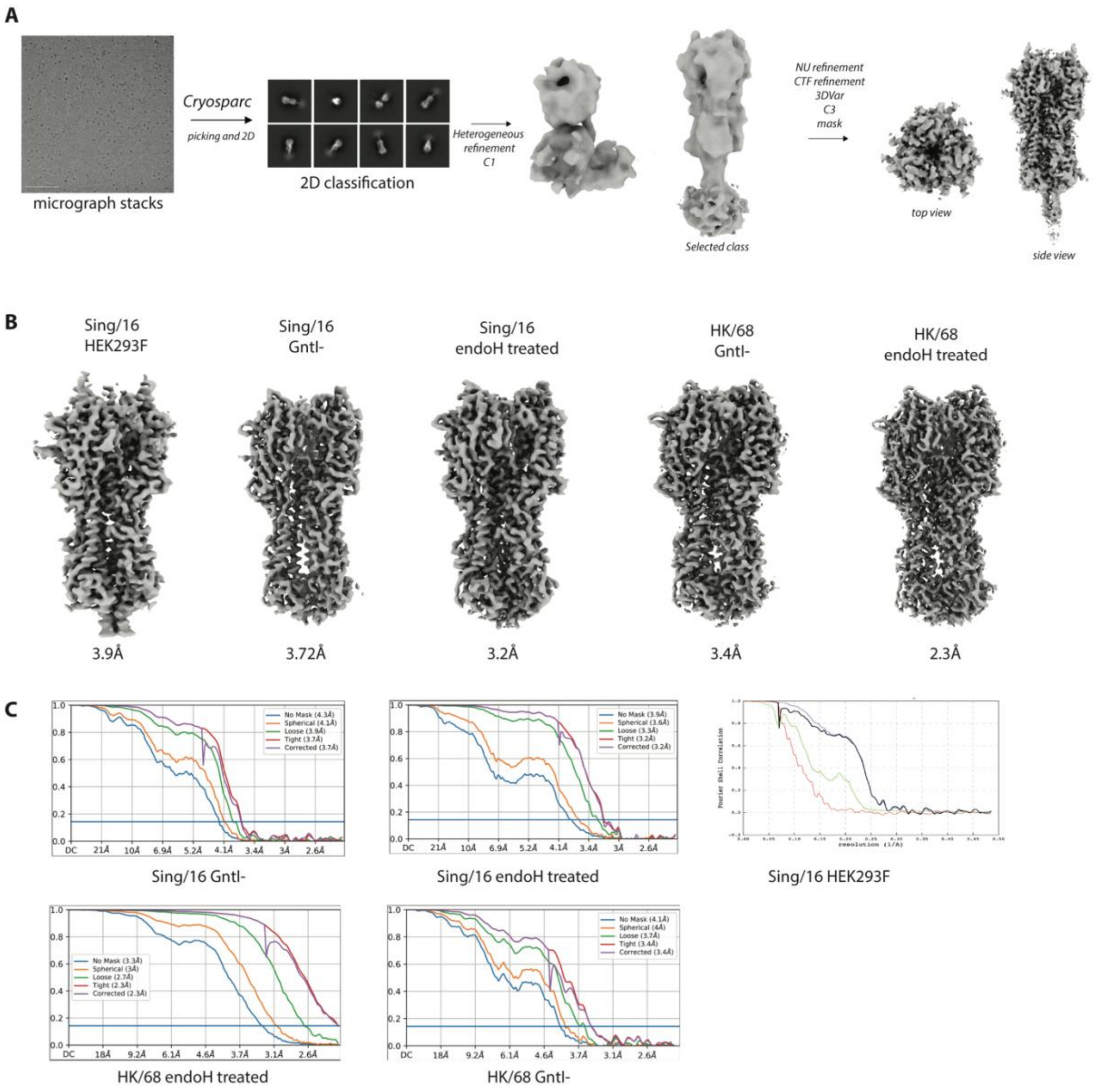

**Supplementary Figure 3. Cryo-EM processing workflow**. (**A**) Schematic representation of generic cryo-EM data processing workflow. (**B**) Individual cryo-EM maps. (**C**) Fourier shell correlation (FSC) plots corresponding the reconstructions shown in panel B.

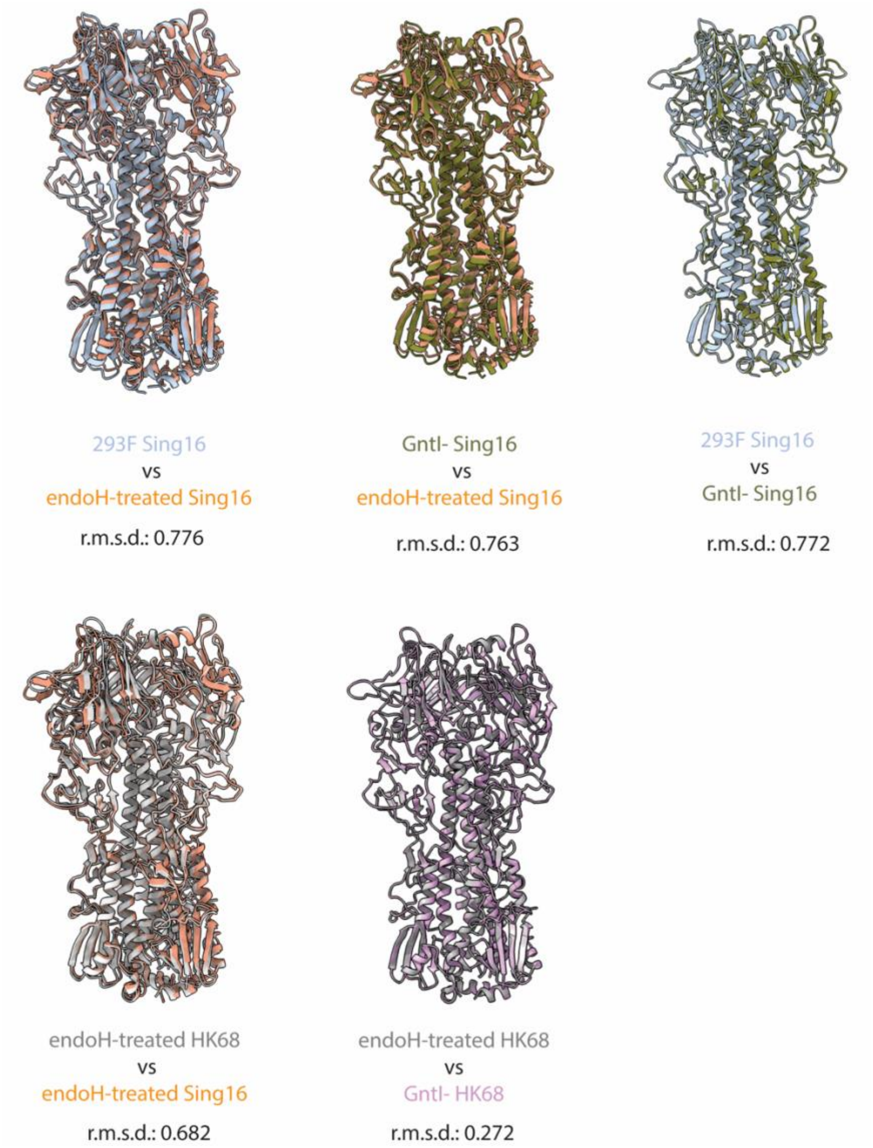

**Supplementary Figure 4. Overlay of cryo-EM models of Sing/16 and HK/68 expressed in different cell lines and after Endo H treated.** Cryo-EM models were overlayed and pairwise r.m.s.d. was performed with one point per residue using ChimeraX.

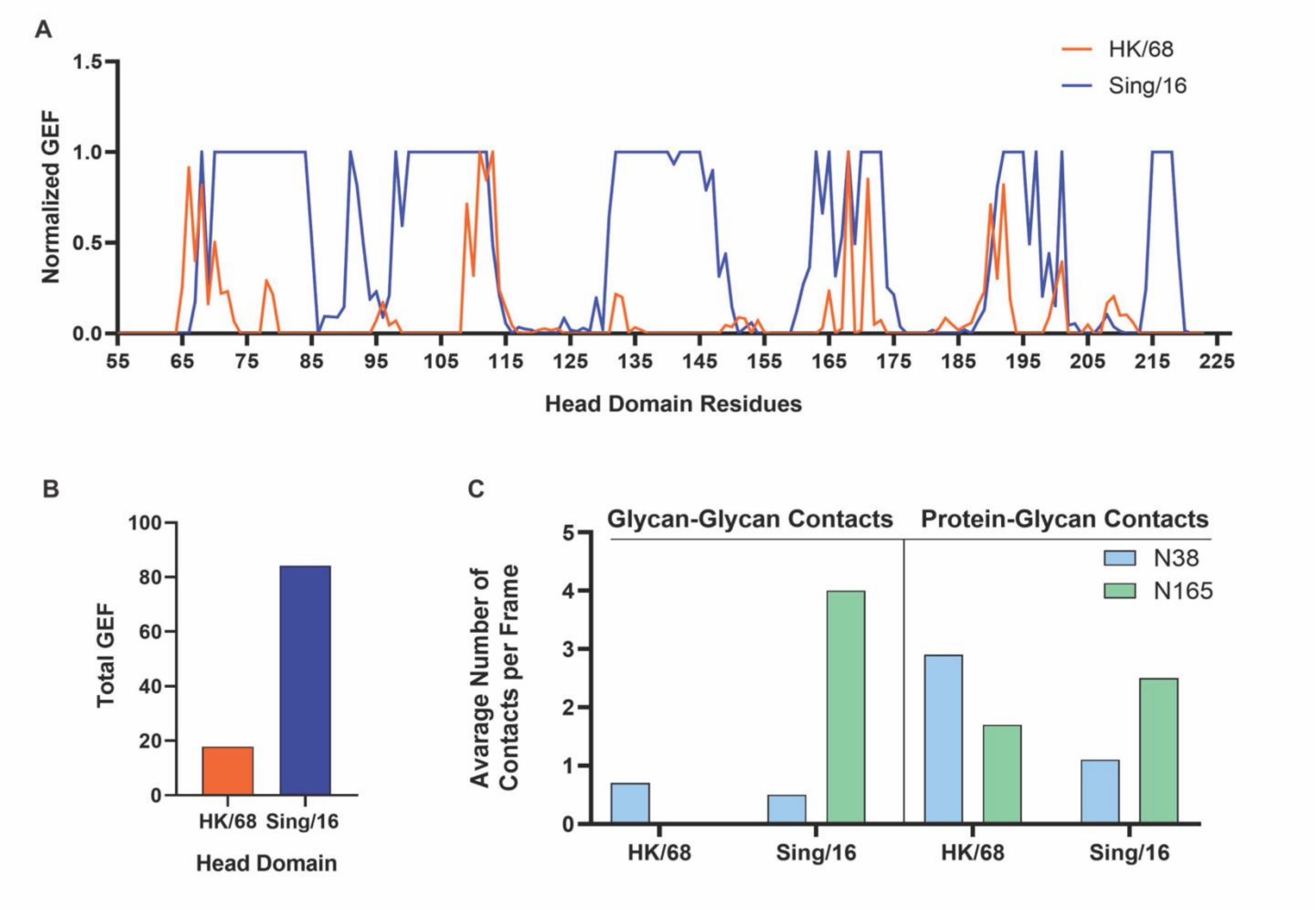

**Supplementary Figure 5. Glycan Shield Analysis**. **(A)** Residue-wise average normalized glycan encounter factor (GEF), which is a quantification of the glycan shielding effect over the HA head domain residues of C52 to C277. HK/68 residues shown in red, and Sing/16 in blue. **(B)** Sum of normalized GEF over all residues of the HA head domain **(C)** Glycan-Glycan and Glycan-Peptide contacts on the HA head for HK/68 and Sing/16.

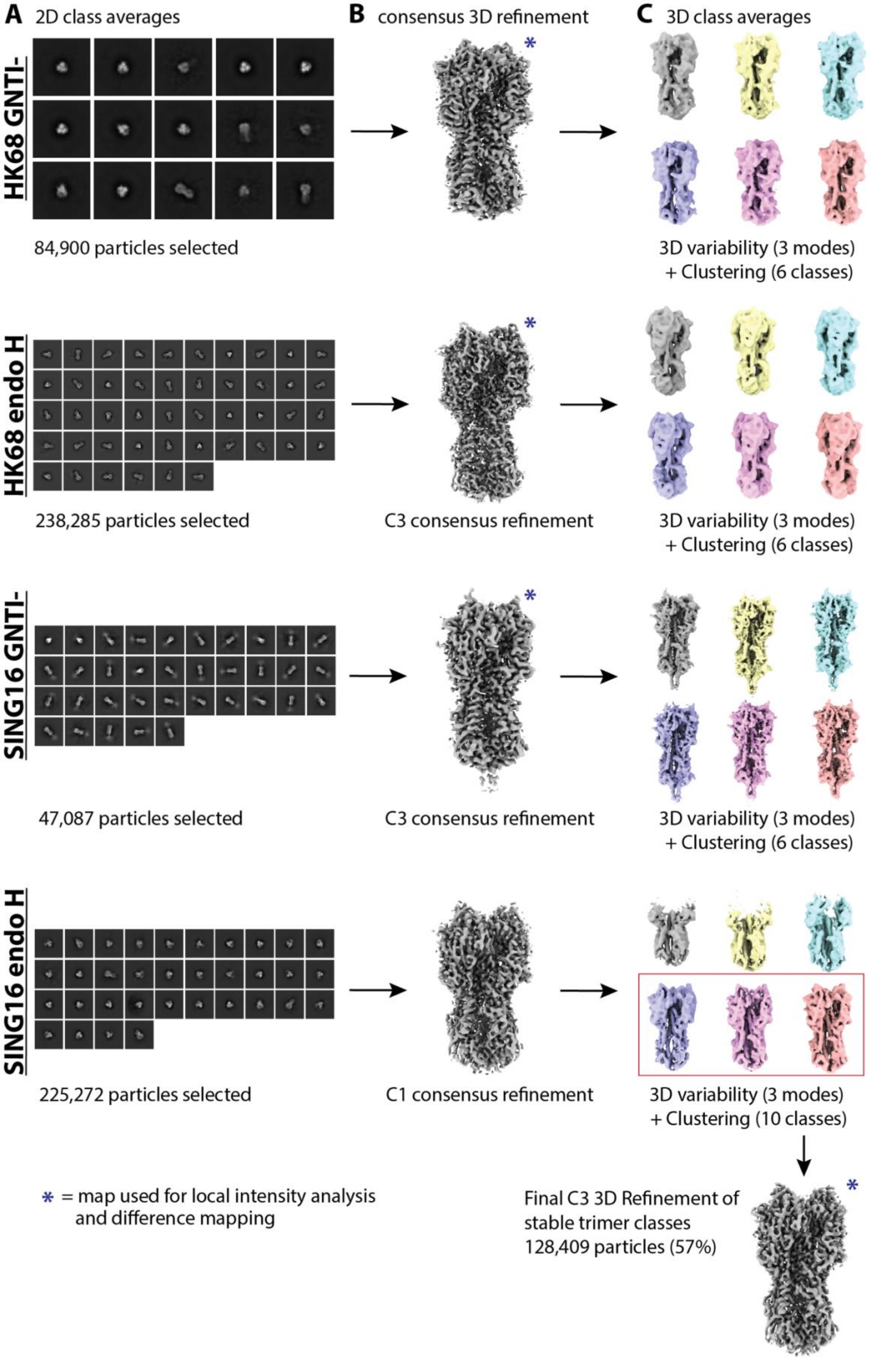

**Supplementary Figure 6. Processing pipeline for discriminating stable and non-intact HA trimers.**

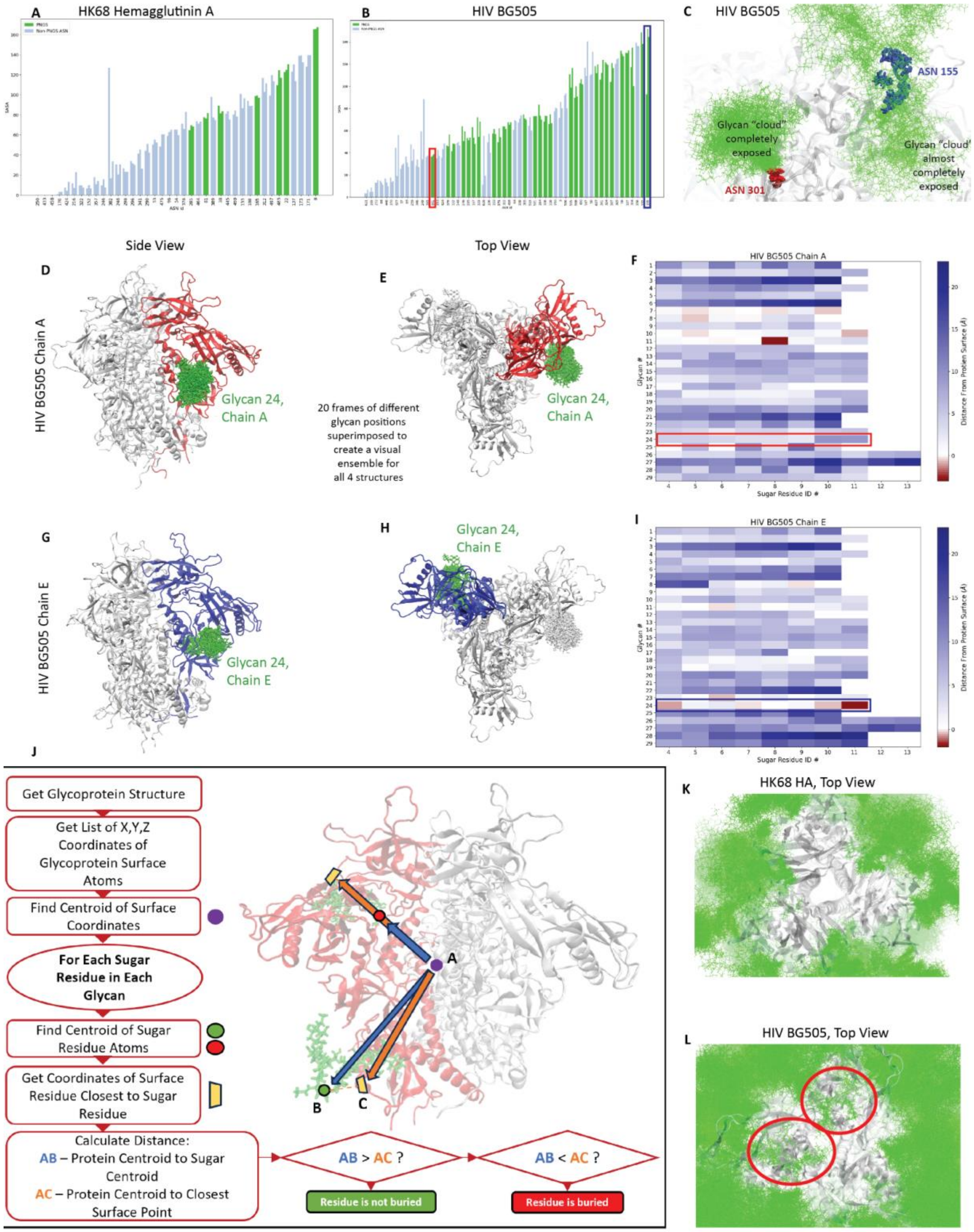

**Supplementary Figure 7. An overview of example outputs of the SASA and Centroid methods of finding buried glycans within viral glycoproteins.** Example plots of SASA vs. the residue identification number of each asparagine in the case of Human Influenza HK68 hemagglutinin A **(A)** and HIV BG505 **(B)** are shown with highlights of the least and greatest SASA PNGS residues shown in red and blue, respectively. A graphical representation of the two SASA extremes of HIV BG505 is shown in **(C)**, where both highlighted residues have similar degrees of glycan exposure regardless of their SASA values. Heat map outputs of the succeeding centroid method in **(F)** and **(I)** depicting the depths below or distance above the protein scaffold surface for each sugar residue of each glycan of an average over 100 frames of an HIV structural trajectory emphasize glycan 24 to demonstrate the visual contrast created between an unburied and buried glycan, respectively. This can be visually seen between the contrast of side and top views of the unburied glycan 24 on HIV BG505 chain A in **(D)** and **(E)**, and the buried glycan 24 on HIV BG505 chain E in **(G)** and **(H)**. The decision tree and graphical depiction in **(J)** shows the logic behind the centroidal method of identifying buried glycans. Finally, two top-down renders of 50 frames of all glycans of **(K)** (Human Influenza HK68 Hemagglutinin A) and **(L)** (HIV) display the contrast between the degree of glycan burial between a structure with almost no glycan burial and one with a high degree of glycan burial highlighted in orange, respectively.

**Supplementary Table 1. Number and percentage of circulating strains containing NXT/S motif at certain amino acid positions**

| Number | 1 | 2 | 3 | 4 | 5 | 6 | 7 | 8 | 9 | 10 | 11 | 12 | 13 | 14 | 15 | 16 | 17 | Total of criculating strains |
|---|---|---|---|---|---|---|---|---|---|---|---|---|---|---|---|---|---|---|
| Position | 6 | 7 | 8 | 22 | 38 | 45 | 63 | 81 | 122 | 126 | 133 | 144 | 158 | 165 | 246 | 276 | 483 | |
| Year | | | | | | | | | | | | | | | | | | |
| 1968 | 0 | 0 | 22 | 22 | 22 | 0 | 5 | 17 | 0 | 1 | 0 | 0 | 0 | 22 | 0 | 0 | 21 | 22 |
| 1969 | 0 | 0 | 5 | 5 | 5 | 0 | 2 | 3 | 0 | 0 | 0 | 0 | 0 | 5 | 0 | 0 | 5 | 5 |
| 1970 | 0 | 0 | 4 | 4 | 4 | 0 | 3 | 1 | 0 | 0 | 0 | 0 | 0 | 4 | 0 | 0 | 4 | 4 |
| 1971 | 0 | 0 | 4 | 8 | 8 | 0 | 0 | 8 | 0 | 0 | 0 | 0 | 0 | 8 | 0 | 0 | 8 | 8 |
| 1972 | 0 | 0 | 12 | 15 | 15 | 0 | 0 | 15 | 0 | 0 | 0 | 0 | 0 | 15 | 0 | 0 | 15 | 15 |
| 1973 | 0 | 0 | 7 | 7 | 7 | 0 | 2 | 7 | 0 | 0 | 0 | 0 | 0 | 7 | 0 | 0 | 7 | 7 |
| 1974 | 0 | 0 | 10 | 10 | 10 | 0 | 6 | 4 | 0 | 7 | 0 | 0 | 0 | 10 | 0 | 0 | 10 | 10 |
| 1975 | 0 | 0 | 10 | 10 | 9 | 0 | 6 | 4 | 0 | 5 | 0 | 0 | 0 | 10 | 0 | 0 | 10 | 10 |
| 1976 | 0 | 0 | 12 | 12 | 12 | 0 | 10 | 2 | 0 | 10 | 0 | 0 | 0 | 11 | 0 | 0 | 12 | 12 |
| 1977 | 0 | 0 | 11 | 11 | 11 | 0 | 9 | 2 | 0 | 9 | 0 | 0 | 0 | 11 | 0 | 0 | 11 | 11 |
| 1978 | 0 | 0 | 7 | 7 | 7 | 0 | 6 | 1 | 0 | 6 | 0 | 0 | 0 | 7 | 0 | 0 | 7 | 7 |
| 1979 | 0 | 0 | 4 | 4 | 4 | 0 | 3 | 1 | 0 | 3 | 0 | 0 | 0 | 4 | 0 | 0 | 4 | 4 |
| 1980 | 0 | 0 | 6 | 8 | 8 | 0 | 7 | 1 | 3 | 7 | 0 | 0 | 0 | 8 | 2 | 0 | 8 | 8 |
| 1981 | 0 | 0 | 3 | 3 | 3 | 0 | 2 | 1 | 2 | 2 | 0 | 0 | 0 | 3 | 0 | 0 | 3 | 3 |
| 1982 | 0 | 0 | 10 | 10 | 10 | 0 | 9 | 1 | 1 | 9 | 0 | 3 | 0 | 10 | 7 | 0 | 10 | 10 |
| 1983 | 0 | 0 | 7 | 7 | 7 | 0 | 6 | 1 | 1 | 6 | 0 | 0 | 0 | 7 | 4 | 0 | 7 | 7 |
| 1984 | 0 | 0 | 5 | 5 | 5 | 0 | 4 | 1 | 0 | 4 | 0 | 0 | 0 | 5 | 4 | 0 | 5 | 5 |
| 1985 | 0 | 0 | 14 | 14 | 14 | 0 | 13 | 1 | 0 | 13 | 0 | 0 | 0 | 14 | 13 | 0 | 14 | 14 |
| 1986 | 0 | 0 | 8 | 8 | 8 | 0 | 7 | 1 | 0 | 7 | 0 | 0 | 0 | 8 | 7 | 0 | 8 | 8 |
| 1987 | 0 | 0 | 6 | 6 | 6 | 0 | 5 | 1 | 0 | 4 | 0 | 0 | 0 | 6 | 4 | 0 | 6 | 6 |
| 1988 | 0 | 0 | 11 | 11 | 11 | 0 | 10 | 1 | 0 | 10 | 0 | 0 | 0 | 11 | 10 | 0 | 11 | 11 |
| 1989 | 0 | 0 | 20 | 20 | 20 | 1 | 19 | 1 | 0 | 18 | 0 | 0 | 0 | 20 | 19 | 0 | 20 | 20 |
| 1990 | 0 | 0 | 9 | 9 | 9 | 2 | 8 | 1 | 0 | 8 | 0 | 0 | 0 | 9 | 8 | 0 | 9 | 9 |
| 1991 | 0 | 0 | 20 | 20 | 20 | 3 | 19 | 1 | 0 | 19 | 0 | 0 | 0 | 20 | 19 | 0 | 20 | 20 |
| 1992 | 0 | 0 | 21 | 21 | 21 | 0 | 20 | 1 | 0 | 20 | 0 | 0 | 0 | 21 | 19 | 4 | 21 | 21 |
| 1993 | 0 | 0 | 68 | 68 | 68 | 0 | 67 | 1 | 0 | 67 | 0 | 0 | 0 | 68 | 59 | 54 | 68 | 68 |
| 1994 | 0 | 0 | 45 | 45 | 45 | 0 | 44 | 1 | 0 | 44 | 0 | 0 | 0 | 45 | 39 | 42 | 45 | 45 |
| 1995 | 0 | 0 | 46 | 46 | 46 | 0 | 45 | 1 | 0 | 43 | 0 | 0 | 0 | 45 | 38 | 10 | 46 | 46 |
| 1996 | 0 | 0 | 56 | 55 | 56 | 3 | 55 | 1 | 18 | 53 | 18 | 0 | 0 | 56 | 50 | 3 | 56 | 56 |
| 1997 | 0 | 0 | 38 | 39 | 39 | 1 | 38 | 1 | 18 | 37 | 20 | 0 | 0 | 38 | 33 | 3 | 39 | 39 |
| 1998 | 1 | 0 | 43 | 44 | 44 | 0 | 43 | 1 | 38 | 43 | 34 | 6 | 0 | 44 | 43 | 0 | 44 | 44 |
| 1999 | 0 | 0 | 111 | 112 | 111 | 3 | 111 | 1 | 108 | 111 | 111 | 42 | 0 | 109 | 108 | 0 | 112 | 112 |
| 2000 | 0 | 0 | 85 | 86 | 86 | 0 | 85 | 1 | 85 | 85 | 85 | 37 | 0 | 83 | 84 | 0 | 86 | 86 |
| 2001 | 0 | 0 | 50 | 51 | 51 | 2 | 50 | 1 | 49 | 48 | 50 | 29 | 0 | 50 | 50 | 0 | 51 | 51 |
| 2002 | 0 | 0 | 110 | 115 | 115 | 4 | 114 | 1 | 114 | 113 | 114 | 22 | 0 | 112 | 114 | 0 | 115 | 115 |
| 2003 | 0 | 1 | 184 | 187 | 187 | 4 | 186 | 1 | 182 | 89 | 186 | 172 | 0 | 182 | 185 | 0 | 187 | 187 |
| 2004 | 0 | 0 | 149 | 152 | 152 | 2 | 150 | 1 | 148 | 130 | 151 | 140 | 0 | 150 | 148 | 0 | 152 | 152 |
| 2005 | 0 | 0 | 151 | 154 | 154 | 3 | 153 | 1 | 145 | 151 | 152 | 148 | 0 | 152 | 150 | 0 | 154 | 154 |
| 2006 | 0 | 0 | 85 | 85 | 85 | 1 | 82 | 1 | 82 | 80 | 84 | 80 | 0 | 85 | 84 | 0 | 85 | 85 |
| 2007 | 1 | 1 | 228 | 232 | 232 | 25 | 231 | 1 | 222 | 218 | 229 | 212 | 0 | 229 | 230 | 0 | 232 | 232 |
| 2008 | 0 | 0 | 169 | 171 | 171 | 1 | 169 | 1 | 166 | 166 | 170 | 168 | 0 | 170 | 169 | 0 | 171 | 171 |
| 2009 | 0 | 0 | 296 | 301 | 299 | 5 | 299 | 1 | 293 | 297 | 289 | 140 | 0 | 298 | 297 | 0 | 301 | 301 |
| 2010 | 1 | 0 | 301 | 310 | 309 | 54 | 309 | 1 | 292 | 306 | 299 | 275 | 0 | 309 | 306 | 0 | 291 | 310 |
| 2011 | 1 | 0 | 432 | 452 | 452 | 123 | 449 | 1 | 426 | 447 | 434 | 409 | 0 | 450 | 444 | 0 | 451 | 452 |
| 2012 | 1 | 0 | 595 | 641 | 643 | 347 | 640 | 1 | 621 | 569 | 636 | 598 | 0 | 639 | 628 | 0 | 641 | 643 |
| 2013 | 2 | 1 | 486 | 514 | 517 | 478 | 515 | 1 | 499 | 261 | 513 | 495 | 4 | 516 | 512 | 0 | 517 | 517 |
| 2014 | 1 | 1 | 634 | 644 | 643 | 636 | 643 | 1 | 550 | 376 | 632 | 310 | 271 | 643 | 636 | 0 | 641 | 644 |
| 2015 | 0 | 2 | 865 | 872 | 877 | 869 | 873 | 1 | 798 | 759 | 861 | 143 | 613 | 875 | 863 | 0 | 876 | 877 |
| 2016 | 2 | 1 | 819 | 846 | 850 | 838 | 848 | 1 | 815 | 703 | 798 | 135 | 606 | 849 | 843 | 0 | 850 | 850 |
| 2017 | 0 | 2 | 1308 | 1317 | 1319 | 1311 | 1313 | 1 | 1203 | 1230 | 948 | 15 | 1135 | 1315 | 1304 | 0 | 1316 | 1319 |
| 2018 | 0 | 7 | 841 | 855 | 856 | 849 | 854 | 3 | 830 | 672 | 694 | 3 | 677 | 853 | 855 | 0 | 857 | 857 |
| 2019 | 0 | 1 | 810 | 830 | 834 | 828 | 831 | 1 | 812 | 319 | 722 | 5 | 383 | 833 | 833 | 0 | 832 | 835 |
| 2020 | 0 | 1 | 93 | 92 | 93 | 91 | 92 | 1 | 85 | 31 | 39 | 11 | 73 | 93 | 91 | 0 | 93 | 93 |
| 2021 | 0 | 0 | 167 | 167 | 169 | 167 | 166 | 1 | 164 | 163 | 162 | 0 | 3 | 169 | 167 | 0 | 169 | 169 |
| 2022 | 1 | 9 | 1244 | 1254 | 1258 | 1254 | 1253 | 1 | 1195 | 1247 | 1176 | 6 | 4 | 1254 | 1257 | 0 | 1259 | 1260 |

| Number | 1 | 2 | 3 | 4 | 5 | 6 | 7 | 8 | 9 | 10 | 11 | 12 | 13 | 14 | 15 | 16 | 17 | Total of criculating strains |
|---|---|---|---|---|---|---|---|---|---|---|---|---|---|---|---|---|---|---|
| Position | 6 | 7 | 8 | 22 | 38 | 45 | 63 | 81 | 122 | 126 | 133 | 144 | 158 | 165 | 246 | 276 | 483 | |
| Year | PNGS (%) | | | | | | | | | | | | | | | | | |
| 1968 | 0.0 | 0.0 | 1.0 | 1.0 | 1.0 | 0.0 | 0.2 | 0.8 | 0.0 | 0.0 | 0.0 | 0.0 | 0.0 | 1.0 | 0.0 | 0.0 | 1.0 | 22 |
| 1969 | 0.0 | 0.0 | 1.0 | 1.0 | 1.0 | 0.0 | 0.4 | 0.6 | 0.0 | 0.0 | 0.0 | 0.0 | 0.0 | 1.0 | 0.0 | 0.0 | 1.0 | 5 |
| 1970 | 0.0 | 0.0 | 1.0 | 1.0 | 1.0 | 0.0 | 0.8 | 0.3 | 0.0 | 0.0 | 0.0 | 0.0 | 0.0 | 1.0 | 0.0 | 0.0 | 1.0 | 4 |
| 1971 | 0.0 | 0.0 | 0.5 | 1.0 | 1.0 | 0.0 | 0.0 | 1.0 | 0.0 | 0.0 | 0.0 | 0.0 | 0.0 | 1.0 | 0.0 | 0.0 | 1.0 | 8 |
| 1972 | 0.0 | 0.0 | 0.8 | 1.0 | 1.0 | 0.0 | 0.0 | 1.0 | 0.0 | 0.0 | 0.0 | 0.0 | 0.0 | 1.0 | 0.0 | 0.0 | 1.0 | 15 |
| 1973 | 0.0 | 0.0 | 1.0 | 1.0 | 1.0 | 0.0 | 0.3 | 1.0 | 0.0 | 0.0 | 0.0 | 0.0 | 0.0 | 1.0 | 0.0 | 0.0 | 1.0 | 7 |
| 1974 | 0.0 | 0.0 | 1.0 | 1.0 | 1.0 | 0.0 | 0.6 | 0.4 | 0.0 | 0.7 | 0.0 | 0.0 | 0.0 | 1.0 | 0.0 | 0.0 | 1.0 | 10 |
| 1975 | 0.0 | 0.0 | 1.0 | 1.0 | 0.9 | 0.0 | 0.6 | 0.4 | 0.0 | 0.5 | 0.0 | 0.0 | 0.0 | 1.0 | 0.0 | 0.0 | 1.0 | 10 |
| 1976 | 0.0 | 0.0 | 1.0 | 1.0 | 1.0 | 0.0 | 0.8 | 0.2 | 0.0 | 0.8 | 0.0 | 0.0 | 0.0 | 0.9 | 0.0 | 0.0 | 1.0 | 12 |
| 1977 | 0.0 | 0.0 | 1.0 | 1.0 | 1.0 | 0.0 | 0.8 | 0.2 | 0.0 | 0.8 | 0.0 | 0.0 | 0.0 | 1.0 | 0.0 | 0.0 | 1.0 | 11 |
| 1978 | 0.0 | 0.0 | 1.0 | 1.0 | 1.0 | 0.0 | 0.9 | 0.1 | 0.0 | 0.9 | 0.0 | 0.0 | 0.0 | 1.0 | 0.0 | 0.0 | 1.0 | 7 |
| 1979 | 0.0 | 0.0 | 1.0 | 1.0 | 1.0 | 0.0 | 0.8 | 0.3 | 0.0 | 0.8 | 0.0 | 0.0 | 0.0 | 1.0 | 0.0 | 0.0 | 1.0 | 4 |
| 1980 | 0.0 | 0.0 | 0.8 | 1.0 | 1.0 | 0.0 | 0.9 | 0.1 | 0.4 | 0.9 | 0.0 | 0.0 | 0.0 | 1.0 | 0.3 | 0.0 | 1.0 | 8 |
| 1981 | 0.0 | 0.0 | 1.0 | 1.0 | 1.0 | 0.0 | 0.7 | 0.3 | 0.7 | 0.7 | 0.0 | 0.0 | 0.0 | 1.0 | 0.0 | 0.0 | 1.0 | 3 |
| 1982 | 0.0 | 0.0 | 1.0 | 1.0 | 1.0 | 0.0 | 0.9 | 0.1 | 0.1 | 0.9 | 0.0 | 0.3 | 0.0 | 1.0 | 0.7 | 0.0 | 1.0 | 10 |
| 1983 | 0.0 | 0.0 | 1.0 | 1.0 | 1.0 | 0.0 | 0.9 | 0.1 | 0.1 | 0.9 | 0.0 | 0.0 | 0.0 | 1.0 | 0.6 | 0.0 | 1.0 | 7 |
| 1984 | 0.0 | 0.0 | 1.0 | 1.0 | 1.0 | 0.0 | 0.8 | 0.2 | 0.0 | 0.8 | 0.0 | 0.0 | 0.0 | 1.0 | 0.8 | 0.0 | 1.0 | 5 |
| 1985 | 0.0 | 0.0 | 1.0 | 1.0 | 1.0 | 0.0 | 0.9 | 0.1 | 0.0 | 0.9 | 0.0 | 0.0 | 0.0 | 1.0 | 0.9 | 0.0 | 1.0 | 14 |
| 1986 | 0.0 | 0.0 | 1.0 | 1.0 | 1.0 | 0.0 | 0.9 | 0.1 | 0.0 | 0.9 | 0.0 | 0.0 | 0.0 | 1.0 | 0.9 | 0.0 | 1.0 | 8 |
| 1987 | 0.0 | 0.0 | 1.0 | 1.0 | 1.0 | 0.0 | 0.8 | 0.2 | 0.0 | 0.7 | 0.0 | 0.0 | 0.0 | 1.0 | 0.7 | 0.0 | 1.0 | 6 |
| 1988 | 0.0 | 0.0 | 1.0 | 1.0 | 1.0 | 0.0 | 0.9 | 0.1 | 0.0 | 0.9 | 0.0 | 0.0 | 0.0 | 1.0 | 0.9 | 0.0 | 1.0 | 11 |
| 1989 | 0.0 | 0.0 | 1.0 | 1.0 | 1.0 | 0.1 | 1.0 | 0.1 | 0.0 | 0.9 | 0.0 | 0.0 | 0.0 | 1.0 | 1.0 | 0.0 | 1.0 | 20 |
| 1990 | 0.0 | 0.0 | 1.0 | 1.0 | 1.0 | 0.2 | 0.9 | 0.1 | 0.0 | 0.9 | 0.0 | 0.0 | 0.0 | 1.0 | 0.9 | 0.0 | 1.0 | 9 |
| 1991 | 0.0 | 0.0 | 1.0 | 1.0 | 1.0 | 0.2 | 1.0 | 0.1 | 0.0 | 1.0 | 0.0 | 0.0 | 0.0 | 1.0 | 1.0 | 0.0 | 1.0 | 20 |
| 1992 | 0.0 | 0.0 | 1.0 | 1.0 | 1.0 | 0.0 | 1.0 | 0.0 | 0.0 | 1.0 | 0.0 | 0.0 | 0.0 | 1.0 | 0.9 | 0.2 | 1.0 | 21 |
| 1993 | 0.0 | 0.0 | 1.0 | 1.0 | 1.0 | 0.0 | 1.0 | 0.0 | 0.0 | 1.0 | 0.0 | 0.0 | 0.0 | 1.0 | 0.9 | 0.8 | 1.0 | 68 |
| 1994 | 0.0 | 0.0 | 1.0 | 1.0 | 1.0 | 0.0 | 1.0 | 0.0 | 0.0 | 1.0 | 0.0 | 0.0 | 0.0 | 1.0 | 0.9 | 0.9 | 1.0 | 45 |
| 1995 | 0.0 | 0.0 | 1.0 | 1.0 | 1.0 | 0.0 | 1.0 | 0.0 | 0.0 | 0.9 | 0.0 | 0.0 | 0.0 | 1.0 | 0.8 | 0.2 | 1.0 | 46 |
| 1996 | 0.0 | 0.0 | 1.0 | 1.0 | 1.0 | 0.1 | 1.0 | 0.0 | 0.3 | 0.9 | 0.3 | 0.0 | 0.0 | 1.0 | 0.9 | 0.1 | 1.0 | 56 |
| 1997 | 0.0 | 0.0 | 1.0 | 1.0 | 1.0 | 0.0 | 1.0 | 0.0 | 0.5 | 0.9 | 0.5 | 0.0 | 0.0 | 1.0 | 0.8 | 0.1 | 1.0 | 39 |
| 1998 | 0.0 | 0.0 | 1.0 | 1.0 | 1.0 | 0.0 | 1.0 | 0.0 | 0.9 | 1.0 | 0.8 | 0.1 | 0.0 | 1.0 | 1.0 | 0.0 | 1.0 | 44 |
| 1999 | 0.0 | 0.0 | 1.0 | 1.0 | 1.0 | 0.0 | 1.0 | 0.0 | 1.0 | 1.0 | 1.0 | 0.4 | 0.0 | 1.0 | 1.0 | 0.0 | 1.0 | 112 |
| 2000 | 0.0 | 0.0 | 1.0 | 1.0 | 1.0 | 0.0 | 1.0 | 0.0 | 1.0 | 1.0 | 1.0 | 0.4 | 0.0 | 1.0 | 1.0 | 0.0 | 1.0 | 86 |
| 2001 | 0.0 | 0.0 | 1.0 | 1.0 | 1.0 | 0.0 | 1.0 | 0.0 | 1.0 | 0.9 | 1.0 | 0.6 | 0.0 | 1.0 | 1.0 | 0.0 | 1.0 | 51 |
| 2002 | 0.0 | 0.0 | 1.0 | 1.0 | 1.0 | 0.0 | 1.0 | 0.0 | 1.0 | 1.0 | 1.0 | 0.2 | 0.0 | 1.0 | 1.0 | 0.0 | 1.0 | 115 |
| 2003 | 0.0 | 0.0 | 1.0 | 1.0 | 1.0 | 0.0 | 1.0 | 0.0 | 1.0 | 0.5 | 1.0 | 0.9 | 0.0 | 1.0 | 1.0 | 0.0 | 1.0 | 187 |
| 2004 | 0.0 | 0.0 | 1.0 | 1.0 | 1.0 | 0.0 | 1.0 | 0.0 | 1.0 | 0.9 | 1.0 | 0.9 | 0.0 | 1.0 | 1.0 | 0.0 | 1.0 | 152 |
| 2005 | 0.0 | 0.0 | 1.0 | 1.0 | 1.0 | 0.0 | 1.0 | 0.0 | 0.9 | 1.0 | 1.0 | 1.0 | 0.0 | 1.0 | 1.0 | 0.0 | 1.0 | 154 |
| 2006 | 0.0 | 0.0 | 1.0 | 1.0 | 1.0 | 0.0 | 1.0 | 0.0 | 1.0 | 0.9 | 1.0 | 0.9 | 0.0 | 1.0 | 1.0 | 0.0 | 1.0 | 85 |
| 2007 | 0.0 | 0.0 | 1.0 | 1.0 | 1.0 | 0.1 | 1.0 | 0.0 | 1.0 | 0.9 | 1.0 | 0.9 | 0.0 | 1.0 | 1.0 | 0.0 | 1.0 | 232 |
| 2008 | 0.0 | 0.0 | 1.0 | 1.0 | 1.0 | 0.0 | 1.0 | 0.0 | 1.0 | 1.0 | 1.0 | 1.0 | 0.0 | 1.0 | 1.0 | 0.0 | 1.0 | 171 |
| 2009 | 0.0 | 0.0 | 1.0 | 1.0 | 1.0 | 0.0 | 1.0 | 0.0 | 1.0 | 1.0 | 1.0 | 0.5 | 0.0 | 1.0 | 1.0 | 0.0 | 1.0 | 301 |
| 2010 | 0.0 | 0.0 | 1.0 | 1.0 | 1.0 | 0.2 | 1.0 | 0.0 | 0.9 | 1.0 | 1.0 | 0.9 | 0.0 | 1.0 | 1.0 | 0.0 | 0.9 | 310 |
| 2011 | 0.0 | 0.0 | 1.0 | 1.0 | 1.0 | 0.3 | 1.0 | 0.0 | 0.9 | 1.0 | 1.0 | 0.9 | 0.0 | 1.0 | 1.0 | 0.0 | 1.0 | 452 |
| 2012 | 0.0 | 0.0 | 0.9 | 1.0 | 1.0 | 0.5 | 1.0 | 0.0 | 1.0 | 0.9 | 1.0 | 0.9 | 0.0 | 1.0 | 1.0 | 0.0 | 1.0 | 643 |
| 2013 | 0.0 | 0.0 | 0.9 | 1.0 | 1.0 | 0.9 | 1.0 | 0.0 | 1.0 | 0.5 | 1.0 | 1.0 | 0.0 | 1.0 | 1.0 | 0.0 | 1.0 | 517 |
| 2014 | 0.0 | 0.0 | 1.0 | 1.0 | 1.0 | 1.0 | 1.0 | 0.0 | 0.9 | 0.6 | 1.0 | 0.5 | 0.4 | 1.0 | 1.0 | 0.0 | 1.0 | 644 |
| 2015 | 0.0 | 0.0 | 1.0 | 1.0 | 1.0 | 1.0 | 1.0 | 0.0 | 0.9 | 0.9 | 1.0 | 0.2 | 0.7 | 1.0 | 1.0 | 0.0 | 1.0 | 877 |
| 2016 | 0.0 | 0.0 | 1.0 | 1.0 | 1.0 | 1.0 | 1.0 | 0.0 | 1.0 | 0.8 | 0.9 | 0.2 | 0.7 | 1.0 | 1.0 | 0.0 | 1.0 | 850 |
| 2017 | 0.0 | 0.0 | 1.0 | 1.0 | 1.0 | 1.0 | 1.0 | 0.0 | 0.9 | 0.9 | 0.7 | 0.0 | 0.9 | 1.0 | 1.0 | 0.0 | 1.0 | 1319 |
| 2018 | 0.0 | 0.0 | 1.0 | 1.0 | 1.0 | 1.0 | 1.0 | 0.0 | 1.0 | 0.8 | 0.8 | 0.0 | 0.8 | 1.0 | 1.0 | 0.0 | 1.0 | 857 |
| 2019 | 0.0 | 0.0 | 1.0 | 1.0 | 1.0 | 1.0 | 1.0 | 0.0 | 1.0 | 0.4 | 0.9 | 0.0 | 0.5 | 1.0 | 1.0 | 0.0 | 1.0 | 835 |
| 2020 | 0.0 | 0.0 | 1.0 | 1.0 | 1.0 | 1.0 | 1.0 | 0.0 | 0.9 | 0.3 | 0.4 | 0.1 | 0.8 | 1.0 | 1.0 | 0.0 | 1.0 | 93 |
| 2021 | 0.0 | 0.0 | 1.0 | 1.0 | 1.0 | 1.0 | 1.0 | 0.0 | 1.0 | 1.0 | 1.0 | 0.0 | 0.0 | 1.0 | 1.0 | 0.0 | 1.0 | 169 |
| 2022 | 0.0 | 0.0 | 1.0 | 1.0 | 1.0 | 1.0 | 1.0 | 0.0 | 0.9 | 1.0 | 0.9 | 0.0 | 0.0 | 1.0 | 1.0 | 0.0 | 1.0 | 1260 |

**Supplementary Table 2. Hemagglutination inhibition assay using sera dilutions from ferrets vaccinated with H3 from 1968-2016.**

| Sera treated with turkey erythrocytes<br>**Start serum dilution is 1/20**<br>Hi in **U-bottom** and 20nM Oseltamivir (end conc.) | | | | | | |
|---|---|---|---|---|---|---|
| | | | Passage | A/Bilth/16190/68 | A/Neth/312/03 | A/Sing/INFIMH-16-0019/2016 |
| | HA | VD | | F10051 | F04001 | F17042 |
| A/Bilthoven/16190/1968 | 128 | 4 | tMK5MDCK2 10-2 | **2560** | <10 | <10 |
| NIB-104 Reass A/Singapore/INFIMH-16-0019/16 | 128 | 4 | E6 11-Sep-2017 | <10 | **10240** | <10 |
| A/Neth/312/2003 | 128 | 4 | xMDCK1 25-Jun-2010 | <10 | <10 | **160** |
| Serum control | | | | <10 | <10 | <10 |

### Supplementary Table 3.

#### Cryo-EM data collection, refinement and validation statistics

| | HA A/Hong kong/1/68 produced in GntI- cells<br><br>(EMDB-45997)<br>(PDB 9CXT) | EndoH-treated HA A/Hong kong/1/68<br><br>(EMDB-45998)<br>(PDB 9CXU) | HA A/Sing/INFIMH/16 produced in 293F cells<br><br>(EMDB-46500)<br>(PDB 9D2M) | HA A/Sing/INFIMH/16 produced in GntI- cells<br><br>(EMDB-46477)<br>(PDB 9D1U) | EndoH-treated HA A/Sing/INFIMH/16<br><br>(EMDB-46466)<br>(PDB 9D0Y) |
|---|---|---|---|---|---|
| **Data collection and processing** | | | | | |
| Microscope | Titan Krios | Arctica | Titan Krios | Arctica | Arctica |
| Magnification | 29,000x | 36,000x | 29,000x | 36,000 | 36,000 |
| Voltage (kV) | 300 | 200 | 300 | 200 | 200 |
| Electron exposure (e–/Å$^2$) | 50 | 50 | 50 | 50 | 50 |
| Defocus range (μm) | -0.8 to -1.5 | -0.8 to -1.5 | -0.8 to -1.5 | -0.8 to -1.5 | -0.8 to -1.5 |
| Detector | K2 Summit DED | Gatan K2 Summit | K2 Summit DED | Gatan K2 Summit | Gatan K2 Summit |
| Recording mode | Counting | Counting | Counting | Counting | Counting |
| Pixel size (Å) | 1.026 | 1.15 | 1.026 | 1.15 | 1.15 |
| Symmetry imposed | C3 | C3 | C3 | C3 | C3 |
| Micrographs (no.) | 497 | 1,141 | 1,714 | 631 | 1.064 |
| Initial particle images (no.) | 308,888 | 509,765 | 479,174 | 680,190 | 1,313,732 |
| Final particle images (no.) | 59,301 | 332,294 | 76,978 | 58,645 | 52,046 |
| Map resolution (Å) | 3.4 | 2.3 | 3.8 | 3.7 | 3.1 |
| FSC threshold | 0.143 | 0.143 | 0.143 | 0.143 | 0.143 |
| Map sharpening *B* factor (Å$^2$) | -109.1 | -67,0 | -105.69 | -141.4 | -94.8 |
| Map pixel size (Å) | 1.026 | 1.15 | 1.026 | 1.15 | 1.15 |
| Map resolution range (Å) | 2.6-4.2 | 2.3-2.8 | 3.2-4.5 | 3.3-4.2 | 2.3-3.8 |
| **Refinement** | | | | | |
| Initial model used (PDB code) | 9CXT | 9CXU | 9D2M | 9D1U | 9D0Y |
| Model resolution (Å) | 3.4 | 2.3 | 3.8 | 3.7 | 3.1 |
| FSC threshold | 0.5 | 0.5 | 0.5 | 0.5 | 0.5 |
| Model resolution range (Å) | 2.6-4.2 | 2.3-2.8 | 3.2-4.5 | 3.3-4.2 | 2.3-3.8 |
| EMRinger score | 2.36 | 5.49 | 3.31 | 3.82 | 4.56 |
| Model composition | | | | | |
| Non-hydrogen atoms | 11,616 | 11,616 | 12,360 | 12,357 | 11,739 |
| Protein residues | 1,446 | 1,446 | 1,434 | 1,434 | 1,431 |
| Ligands | 18 | 18 | 72 | 72 | 27 |
| Mean *B* factors (Å$^2$) | | | | | |
| Protein | 11.15 | 11.15 | 44.37 | 45.42 | 45.55 |
| Ligand | 33.94 | 33.94 | 61.55 | 96.64 | 70.67 |
| R.m.s. deviations | | | | | |
| Bond lengths (Å) | 0.021 | 0.021 | 0.023 | 0.023 | 0.023 |
| Bond angles (°) | 1.677 | 1.676 | 2.508 | 2.186 | 1.688 |
| Validation | | | | | |
| MolProbity score | 1.02 | 0.98 | 1.44 | 1.22 | 0.81 |
| Clashscore | 2.37 | 2.11 | 5.03 | 4.45 | 1.08 |
| Poor rotamers (%) | 0.00 | 0.00 | 0.00 | 0.00 | 0.00 |
| Ramachandran plot | | | | | |
| Favored (%) | 98.95 | 98.05 | 96.98 | 98.10 | 98.73 |
| Allowed (%) | 1.05 | 1.95 | 3.02 | 1.90 | 1.27 |
| Disallowed (%) | 0.00 | 0.00 | 0.00 | 0.00 | 0.00 |
